# Neutralizing antibodies induced in immunized macaques recognize the CD4-binding site on an occluded-open HIV-1 envelope trimer

**DOI:** 10.1101/2021.11.02.466200

**Authors:** Zhi Yang, Kim-Marie A. Dam, Michael D. Bridges, Magnus A.G. Hoffmann, Andrew T. DeLaitsch, Harry B. Gristick, Amelia Escolano, Rajeev Gautam, Malcolm A. Martin, Michel C. Nussenzweig, Wayne L. Hubbell, Pamela J. Bjorkman

## Abstract

Broadly-neutralizing antibodies (bNAbs) against HIV-1 Env can protect from infection. We characterize Ab1303 and Ab1573, heterologously-neutralizing CD4-binding site (CD4bs) antibodies, isolated from sequentially-immunized macaques. Ab1303/Ab1573 binding is observed only when Env trimers are not constrained in the closed, prefusion conformation. Fab-Env cryo-EM structures show that both antibodies recognize the CD4bs on Env trimer with an ‘occluded-open’ conformation between closed, as targeted by bNAbs, and fully-open, as recognized by CD4. The occluded-open Env trimer conformation includes outwardly-rotated gp120 subunits, but unlike CD4-bound Envs, does not exhibit V1V2 displacement, 4-stranded gp120 bridging sheet, or co-receptor binding site exposure. Inter-protomer distances within trimers measured by double electron-electron resonance spectroscopy suggest an equilibrium between occluded-open and closed Env conformations, consistent with Ab1303/Ab1573 binding stabilizing an existing conformation. Studies of Ab1303/Ab1573 demonstrate that CD4bs neutralizing antibodies that bind open Env trimers can be raised by immunization, thereby informing immunogen design and antibody therapeutic efforts.

## Introduction

Human immunodeficiency virus-1 (HIV-1) is the causative agent behind the ongoing AIDS pandemic affecting millions of people worldwide. Although HIV-1 infection induces neutralizing antibodies against the viral envelope glycoprotein trimer (Env), the large number of viral strains in a single infected person and across the infected population means that commonly-produced strain-specific antibodies do not clear the infection^1^. However, a fraction of infected patients produced broadly neutralizing antibodies (bNAbs) that could provide protection from HIV-1 infection if an efficient means of eliciting such antibodies is developed^2,3^.

Neutralizing antibodies against HIV-1 are exclusively directed against Env, the only viral protein on the surface of the virion^4,5^. HIV-1 Env is a homotrimer of gp120-gp41 heterodimers that mediates fusion of the host and viral membrane bilayers to allow entry of viral RNA into the host cell cytoplasm^6^. Fusion is initiated when the Env gp120 subunit contacts the host receptor CD4, resulting in conformational changes that reveal the binding site for a host coreceptor in the chemokine receptor family^7,8^. Coreceptor binding to gp120 results in further conformational changes including insertion of the gp41 fusion peptide into the host cell membrane^6^.

Conformations of trimeric HIV-1 Envs have been investigated using single-particle cryo-EM to derive structures of soluble, native-like Env trimers lacking membrane and cytoplasmic domains and including stabilizing mutations (SOSIP.664 Envs)^9^. Such structures defined a closed, pre-fusion Env state in which the coreceptor binding site on gp120 variable loop 3 (V3) is shielded by the g120 V1V2 loops^10^, and open CD4-bound Env trimer states with outwardly rotated gp120 subunits and V1V2 loops displaced by ~40Å to expose the V3 loops and coreceptor binding site^11–14^.

Structurally-characterized anti-HIV-1 bNAbs recognize the closed, pre-fusion Env state^10^ with the exception of one of the first HIV-1 bNAbs to be discovered: an antibody called b12 that was isolated from a phage display screen^15^. Like more recently-identified bNAbs^16,17^, b12 binds to an epitope overlapping with the CD4-binding site (CD4bs) on gp120^18^. However, the Env trimer state recognized by b12 represents an ‘occluded-open’ conformation in which the gp120 subunits are rotated out from the central trimer axis, but V1V2 is not displaced to the sides of the Env trimer^11,12,14^.

As compared with a library screen that would not preserve correct heavy chain-light chain pairing, Ab1303 and Ab1573 were isolated by single cell cloning from SOSIP-binding B cells derived from sequentially-immunized non-human primates (NHPs)^19^. Both antibodies exhibited broad, but weak, heterologous neutralization and were mapped by competition ELISA as recognizing the CD4bs^19^. Here we show that, in common with b12 but not with other CD4bs bNAbs, neither antibody binds Env trimer when it is locked into the closed, prefusion state that is recognized by other CD4bs bNAbs. To elucidate the conformational state of Env recognized by these monoclonal antibodies (mAbs), we solved single-particle cryo-EM structures of a SOSIP Env trimer complexed with either Ab1303 or Ab1573 Fabs. The structures revealed that these mAbs recognized Env trimers with gp120 subunits that had rotated outwards to create an ‘occluded-open’ trimer conformation that differed from the closed, prefusion Env conformation and from the open conformation of CD4-bound Env trimers^11–14^. To further investigate the occluded-open Env trimer conformation, we used double electron-electron resonance (DEER) spectroscopy to determine if this conformation was detectable in a solution of unliganded HIV-1 Env trimers. By measuring inter-protomer distances between V1V2 loops in the presence and absence of Ab1303 and Ab1573, we found evidence for the conformation recognized by these antibodies in unliganded trimers, suggesting that Ab1303 or Ab1573 binding stabilized a pre-existing Env conformation.

In contrast to previous structures of CD4bs bNAb-closed Env trimer complexes^10^ and CD4-bound open Env conformation structures^11–14^, the Ab1303 and Ab1573 structures revealed a new mode of naturally-induced CD4bs antibody-Env interaction. Furthermore, when combined with DEER spectroscopy data, these structures define Env trimer conformational state intermediates between the closed and CD4-bound open conformations. Although Ab1303 and Ab 1573 are not as broad or potent as CD4bs bNAbs isolated from infected human donors, discovery of the Env structure recognized by these mAbs reveals an unanticipated target that could be exploited for immunogen design.

## Results

### Heterologously neutralizing mAbs Ab1303 and Ab1573 were elicited in NHPs after sequential immunization with designed immunogens

The V3-glycan patch immunogen RC1 was modified from the V3 immunogen 11MUTB^20^ by mutating a potential N-linked glycan site (PNGS) to remove the *N*-glycan attached to Asn156_gp120_^21^. RC1 and 11MUTB were both derived from the clade A BG505 SOSIP.664 native-like Env trimer^9^. We constructed RC1-4fill and 11MUTB-4fill by modifying RC1 and 11MUTB, respectively, to insert PNGSs to add glycans to residues 230_gp120_, 241 _gp120_, 289_gp120_, and 344_gp120_ to reduce antibody responses to off-target epitopes^22–24^. Immunogens were multimerized on VLPs using the SpyTag-SpyCatcher system^25,26^ to enhance avidity effects and limit antibody access to the Env trimer base. The mAbs described here were isolated from NHPs immunized sequentially as shown in Figure S1A. As described elsewhere^19^, we obtained weak heterologously neutralizing antisera from the sequentially-immunized NHPs, and mAb sequences were generated by single cell cloning from B cells that were captured as described using BG505 and B41 SOSIP baits^27^. Here we investigated Ab1303 and Ab1573, which unexpectedly recognized the CD4bs rather than the V3-glycan patch that was targeted in the sequential immunization scheme.

Ab1303 sequences were derived from rhesus macaque germline V gene segments *IGHV4-160*01* and *IGLV4-97*01* gene segments for heavy and light chains, respectively, and exhibited 8.3% and 8% amino acid changes due to somatic hypermutations, respectively (Figure S1B). Ab1573 sequences were derived from *IGHV1-198*02* and *IGLV1-64*01* gene segments and contained 7.3% and 10.5% somatic hypermutation changes, respectively (Figure S1B). Neutralizing activities of the two antibodies were reported elsewhere^19^. The HIV-1 strains chosen for neutralization measurements were derived from the 12-strain global panel of HIV-1 reference strains^28^ plus seven other HIV-1 strains including BG505, from which the RC1-4fill and 11MUT-4fill immunogens were derived. We found that Ab1303 neutralized 12 of the 19 cross-clade strain panel with IC_50_ values <100 μg/mL, whereas Ab1573 neutralized five strains in the panel with IC_50_ values <100 μg/mL^19^. Although the neutralization potencies were generally weak, both mAbs exhibited heterologous neutralization, with Ab1303 neutralizing >60% of the viruses in the cross-clade panel when evaluated at high concentrations.

### Ab1303/Ab1573 bound a non-closed Env trimer conformation with varying stoichiometries

To verify that Ab1303 and Ab1573 recognized the CD4bs on Env trimer, we repeated competition ELISA experiments conducted with RC1 trimer^19^, this time using BG505 trimer (Figure 1A). We first immobilized randomly-biotinylated BG505 Env trimers on streptavidin plates and then added antibody Fabs targeting either the CD4bs (3BNC117)^16^, the V3-glycan patch (10-1074)^29^, V1V2 (PG16)^30^, or the fusion peptide (VRC34)^31^. Subsequently, either Ab1303 or Ab1573 IgGs were added, the plates were washed, and the degree of binding was detected. The binding of Ab1303 IgG was essentially unaffected in the Env trimer samples that were complexed with 10-1074, PG16, or VRC34 compared to the control with no competitor (Ab1303 IgG alone), but its binding to BG505 Env was reduced in the presence of 3BNC117 Fab (Figure 1A). Similar results were found for Ab1573, although the presence of PG16 and VRC34 Fabs also reduced the binding somewhat (Figure 1A). From these results, we concluded that both antibodies recognized the CD4bs on BG505 SOSIP.

**Figure 1.**
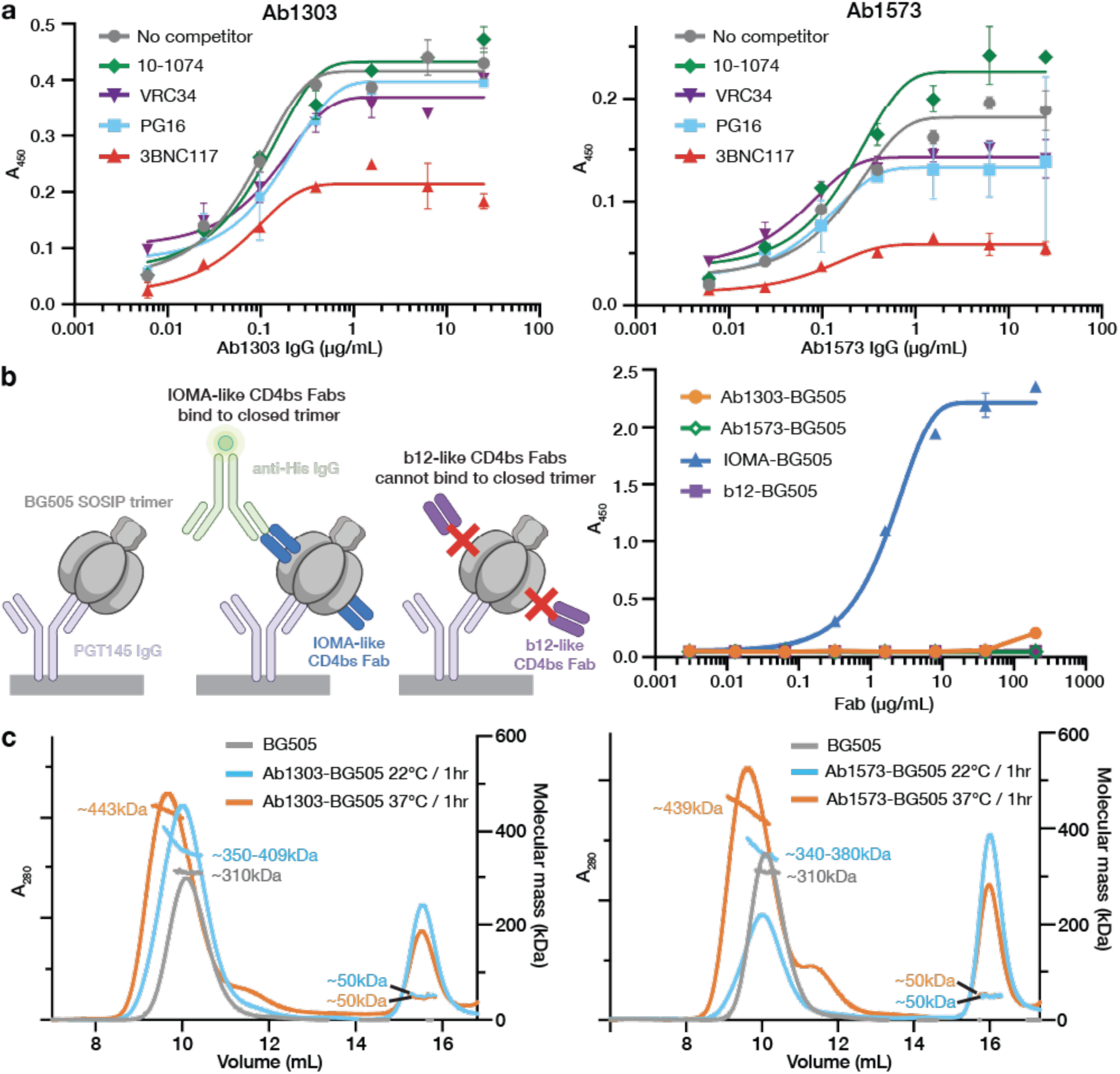
Ab1303 and Ab1573 are CD4bs NAbs that bind an Env conformation other than the closed, prefusion state. (A) Competition ELISA to map the binding sites of Ab1303 (left) and Ab1573 (right) on BG505 SOSIP. Randomly biotinylated BG505 SOSIP trimer was immobilized on a streptavidin plate and then incubated with a Fab from a bNAb that recognizes the V3 loop (10-1074), the fusion peptide (VRC34), the V1V2 loop (PG16), or the CD4bs (3BNC117). Increasing concentrations of Ab1303 or Ab1573 IgG were added to the Fab-BG505 complex, resulting in competition in both cases with the CD4bs bNAb 3BNC117. (B) ELISA to assess binding of Ab1303 and Ab1573 to closed, prefusion conformation of BG505 trimer. Left: schematic of the experiment: PGT145 IgG, which constrains BG505 to a closed prefusion conformation, was immobilized and captured BG505. Binding of His-tagged Fab to BG505 was detected using a labeled anti-His-tag antibody. Right: Ab1303, Ab1573, and b12 do not bind to BG505 captured by PGT145, but IOMA binds to BG505 captured by PGT145. (C) SEC-MALS of Ab1303-BG505 (left) and Ab1573-BG505 (right) complexes incubated at different temperatures. SEC traces were monitored by absorption at 280 nm. Traces and calculated molecular mass distributions for unliganded BG505 (gray) and for Ab1303 Fab or Ab1573 Fab incubated with BG505 SOSIP trimer at 37℃ (orange) or 22°C (blue) for 1 hour are shown.

CD4bs bNAbs such as VRC01, 3BNC117, and IOMA bind closed, prefusion state Env trimers^32–34^. An exception to this finding for CD4bs antibodies is b12, a more weakly neutralizing antibody selected from a phage display derived from antibody genes isolated from an HIV-positive individual bone marrow^15^. Unlike all other human CD4bs bNAbs characterized to date, b12 binds to an “occluded open” Env trimer conformation in which gp120 subunits are rotated outwards from the central trimer axis but the V1V2 loops are not displaced from their positions on top of the gp120 subunits^11^.

To determine if Ab1303 and Ab1573 recognize closed Env trimers, we assessed their ability to bind Env trimer captured with PGT145, a V1V2 bNAb that recognizes a quaternary epitope at the trimer apex^33^. Unlike other V1V2 bNAbs such as PG16 that can bind recognize both closed and open Env trimers^35^, PGT145 locks Envs into a closed, prefusion state^33^. In this experiment, we captured BG505 with PGT145 IgG on an ELISA plate and compared binding of Ab1303, Ab1573, a conventional CD4bs bNAb that recognizes closed Env trimer (IOMA)^32^, and b12 Fab that binds to an “occluded open” trimer (Figure 1B). IOMA showed binding to BG505 captured by PGT145, consistent with IOMA-BG505 complex structure with a closed conformation trimer^32^ (Figure 1B). By contrast, b12 did not bind to BG505 that was captured by PGT145, as the b12 epitope is occluded in the closed Env conformation^11^ (Figure 1B). Similar to the results for b12, Ab1303 and Ab1573 did not bind BG505 that was captured by PGT145, suggesting that the closed Env trimer occludes epitopes for Ab1303 and Ab1573 (Figure 1B).

To further characterized the interactions of Ab1303 and Ab1573 with Env trimer, we performed size-exclusion chromatography coupled with multi-angle light scattering (SEC-MALS) to determine the absolute molecular masses of the complexes and therefore the number of Fabs bound per trimer. Complexes formed by incubating mAb Fabs with BG505 trimer at various temperatures were analyzed by SEC-MALS. Compared to BG505 trimer alone (~310 kDa apparent mass including glycans), incubation of Ab1303 Fab with trimer at 22℃ for one hour resulted in a complex with an average molecular mass of 376 kDa, equivalent to ~1.3 Fabs per trimer, whereas incubation at 37℃ for one hour produced complexes with an average molecular mass of 448 kDa, corresponding to ~3 Fabs per trimer (Figure 1C). In the case of Ab1573 complexes, one-hour incubations at 22℃ and 37℃ produced ~1.2 and ~2.6 copies of Fabs per trimer on average (Figure 1C). Notably, peaks corresponding to the Ab1303 Fab-BG505 complex from the 22℃ incubation condition and the Ab1573 Fab-BG505 complexes from both temperature conditions were broad, consistent with a mixture of sub-stoichiometric populations being present under these conditions. These observations suggest that physiological temperature could result in the antibody binding sites on Env being more accessible, facilitating binding of Ab1303 and Ab1573 by more frequent Env transitions between different conformational states at higher temperature.

### Ab1303 and Ab1573 occlude the CD4bs on gp120

To further explore the interactions of Ab1303 and Ab1573 with Env trimer, we solved 1.51Å and 2.24Å crystal structures of unbound Ab1303 and Ab1573 Fabs (Table S1; Figure S1C) and single-particle cryo-EM structures of BG505 SOSIP.664 complexed with Ab1303 and Ab1573 to resolutions of 4.0Å and 4.1Å, respectively (Figure 2A,B, Figure S2, and Table S2). Prior to cryo-EM data collection, the Fabs were incubated with BG505 at 37℃ for two hours to achieve an approximate 3:1 Fab to BG505 trimer stoichiometry.

**Figure 2.**
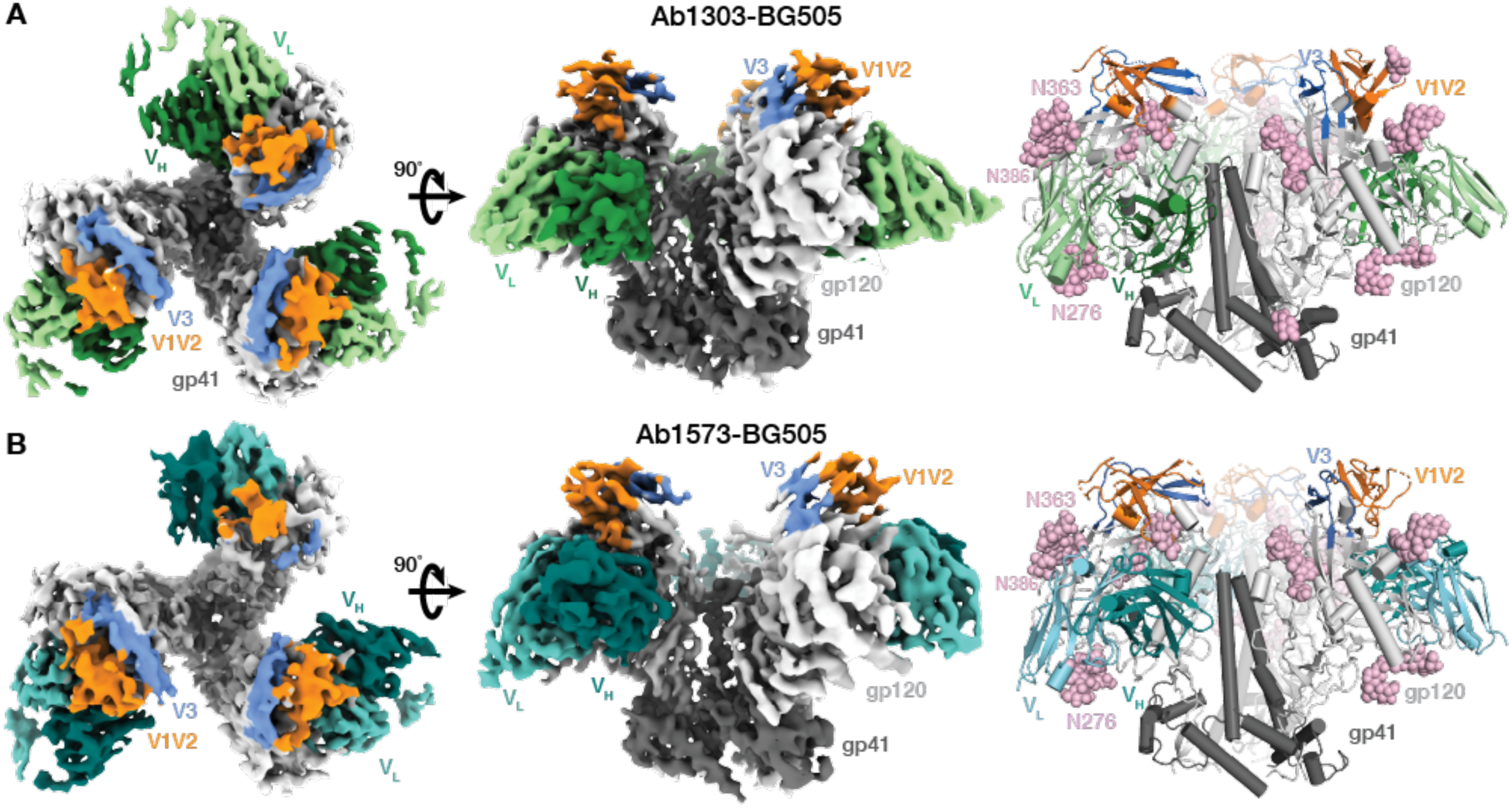
Cryo-EM maps and atomic models of Ab1303-BG505 and Ab1573-BG505 complexes. (A,B) Cryo-EM density maps of Ab1303-BG505 (A) and Ab1573-BG505 (B) complexes shown from the top (left) and side (middle). Right: cartoon diagrams of the structures.

We first compared the structures of the unbound Fabs to their structures when bound to Env trimer. For both antibodies, there were no major structural changes between their free (solved by X-ray crystallography) and bound (solved by cryo-EM) forms: Root mean square deviations, rmsds, for superimposition of free and bound Ab1303 V_H_ and V_L_ (235 Cα atoms) were 0.72Å, and 0.89Å for superimposition of free and bound Ab1573 V_H_ and V_L_ (231 Cα atoms), with minimal differences in the complementarity determining region (CDR) loops (Figure S1D). Thus, Ab1303 and Ab1573 bound their Env antigen targets using preformed antibody combining sites, rather than undergoing structural rearrangements to accommodate their targets.

The cryo-EM structures of both Fab complexes with Env trimer showed three bound Fabs that interacted with the CD4bs of each gp120 protomer (Figure 2A,B; Figure 3). The binding sites for Ab1303 and Ab1573 on gp120 were located in an area that is surrounded by three *N*-glycan patches: the Asn363_gp120_/Asn386_gp120_ glycans located near the base of V3, the Asn197_gp120_ glycan in the V1V2 region, and the Asn276_gp120_ glycan near the bottom of gp120 (Figure 2A,B). The epitope of Ab1303 comprised 1430Å^2^ of buried surface area (BSA) on gp120, of which 879Å^2^ were buried by the heavy chain and 551Å^2^ were buried by the light chain (Figure 3A,G). The heavy chain complementarity determining region 3 (CDRH3) of Ab1303 makes extensive contacts with gp120, including an antibody residue Arg100B_HC_ that is stabilized by neighboring Trp100A_HC_ through cation-π interaction, contributes a salt bridge with gp120 residue Asp457_gp120_ and hydrogen bonds with the Arg456_gp120_ carbonyl group and with the Ser365_gp120_ sidechain, forming a stable interaction network (Figure 3B). Residue Tyr100E_HC_ of CDRH3 hydrogen bonded with Gln428_gp120_ and Asp474_gp120_ (Figure 3B). In the case of Ab1573, 646Å^2^ of surface area was buried by V_H_ and 762Å^2^ buried by VL, composing a total of 1408Å^2^ of BSA on _gp120_ (Figure 3C,G). Two salt bridges were found at the Ab1573-gp120 interface: between CDRH3 residue Asp97_HC_ and Arg476_gp120_, and between CDRH1 residue Arg31_HC_ and Asp113_gp120_ (Figure 3D). Compared with the Ab1573 and Ab1573 footprints on gp120, contacts by b12 are dominated by its V_H_ domain (Figure 3E), with no contacts by V_L_ except for a small region of BSA (340 Å^2^) on V1V2 (Figure 3F,G).

**Figure 3.**
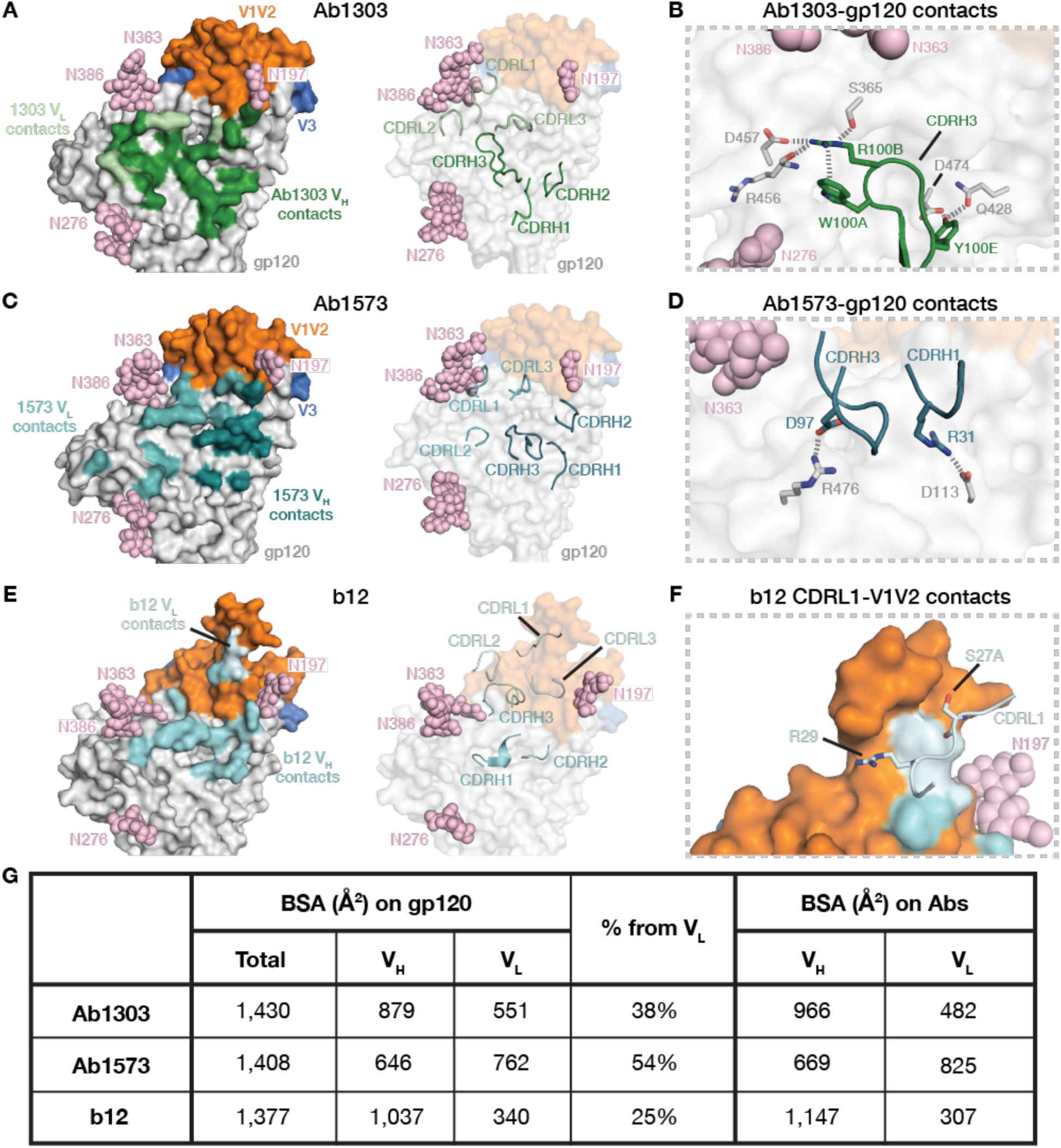
Ab1303 and Ab1573 recognize the CD4bs of BG505 Env trimer. (A) Left: epitope of Ab1303 on gp120 surface with contacts made by antibody heavy and light chains colored in dark and light green, respectively. Right: CDR loops of Ab1303 mapped onto the gp120 surface. (B) Highlighted interactions between gp120 residues and the Ab1303 CDRH3. (C) Left: epitope of Ab1573 on gp120 surface with contacts made by antibody heavy and light chains colored in dark and light teal, respectively. Right: CDR loops of Ab1573 mapped onto the gp120 surface. (D) Highlighted interactions between gp120 residues and the Ab1573 CDRH1 and CDRH3. (E) Left: epitope of b12 on gp120 surface with contacts made by antibody heavy and light chains colored in dark and light cyan, respectively. Right: CDR loops of b12 mapped onto the gp120 surface. (F) Highlighted interactions between gp120 V1V2 and the b12 VL. (G) Table of buried surface areas (BSAs).

In addition to their protein epitopes, Ab1303 and Ab1573 also contacted *N*-linked glycans on gp120. While the Ab1303 light chain was adjacent to the Asn276_gp120_ glycan, the Ab1573 heavy and light chains contacted the N-glycans attached to Asn197_gp120_, Asn276_gp120_, Asn363_gp120_, and Asn386_gp120_, likely resulting in glycan rearrangements to allow binding (Figure 3A,B).

To further characterize the antibody epitope, we compared our structures with those of other CD4bs-Env complexes. The V_H_ domains of Ab1303 and Ab1573 were positioned close to the gp120 inner domain, which is not fully exposed in the closed Env conformation, whereas the V_L_ domains were sandwiched between Asn363_gp120_/Asn386_gp120_ and Asn276_gp120_ glycans (Figure 4). While the b12 V_H_ is positioned similarly on gp120 as the V_H_ domains of Ab1303 and Ab1573 (Figure 3A,C,E), the b12 V_L_ is closer to the gp120 V1V2, which constitutes the only contacts made by the b12 V_L_ with gp120 (Figure 3E,F,G). By contrast, other CD4bs bNAb VHV_L_ domains share similar binding poses; they are located further from the gp120 inner domain and are lined up almost parallel to the trimer three-fold axis, with V_H_ near the Asn363_gp120_/Asn386_gp120_ and Asn197_gp120_ glycans and V_L_ adjacent to the Asn276_gp120_ glycan (Figure 4), and to accommodate such poses, Asn276 glycans need to be displaced further away by antibodies light chains.

**Figure 4.**
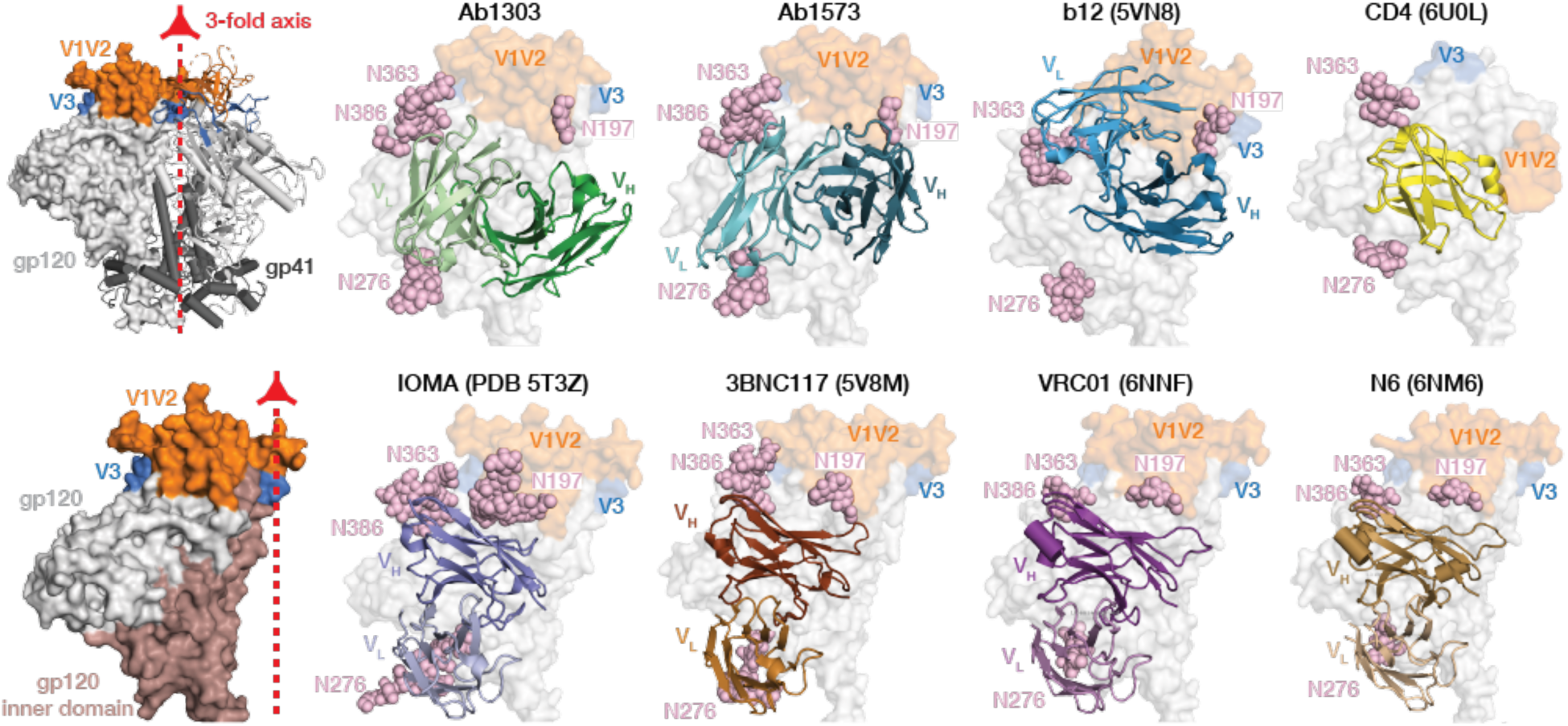
Differences in antibody-binding poses for CD4bs bNAbs compared with CD4. Upper and lower left: surface representation of HIV-1 Env trimer (top) and gp120 monomer (bottom) showing locations of the 3-fold axis relating Env protomers (red arrow), the gp120 inner domain, V1V2, V3, and gp41. Remaining panels: Cartoon diagrams of V_H_-VL domains of the indicated bNAbs bound to HIV-1 gp120 (gray surface, pink spheres for N-linked glycans, V1V2 in orange, V3 in blue). PDB codes: Ab1303 and Ab1573 (this study), b12 (5VN8), CD4 (6U0L), IOMA (5T3Z), 3BNC117 (5V8M), a VRC01 derivative (6NNF), and an N6 derivative (6NM6). The V1V2 and V3 regions were largely disordered in the CD4-bound Env structure (PDB 6U0L).

### Ab1303 and Ab1503 bind an ‘occluded-open’ state of Env trimer

The trimers in the Ab1303-Env and Ab1573-Env complexes differed in conformation from the closed, prefusion Env conformation, each exhibiting a more open state that exposed portions of the gp120 that were otherwise buried (Figure 5A,B). To characterize these differences, we mapped the trimer epitope regions from each antibody-bound open conformation onto a closed, prefusion trimer Env structure. For both complexes, a portion of the epitope was solvent inaccessible in the closed trimer state but was exposed in the antibody-bound open state (Figure 5A,B; red highlighted regions). The Ab1303 contacts that are buried in a closed trimer were contacted exclusively on the occluded, open trimer by its V_H_ domain (Figure 5A), which buried 286Å2 of gp120 surface area that would be inaccessible on a closed trimer. The contact residues buried in the closed Env state by Ab1573 also involved only its V_H_ domain (Figure 5B), burying a discontinuous 137Å2 of gp120 surface area. In addition, docking of Ab1303 (Figure 5C) or Ab1573 (Figure 5D) onto a closed trimer structure results in steric clashes. These results are consistent with the ELISA demonstration that Ab1303 and Ab1573 did not bind the closed, prefusion Env trimers (Figure 1B).

**Figure 5.**
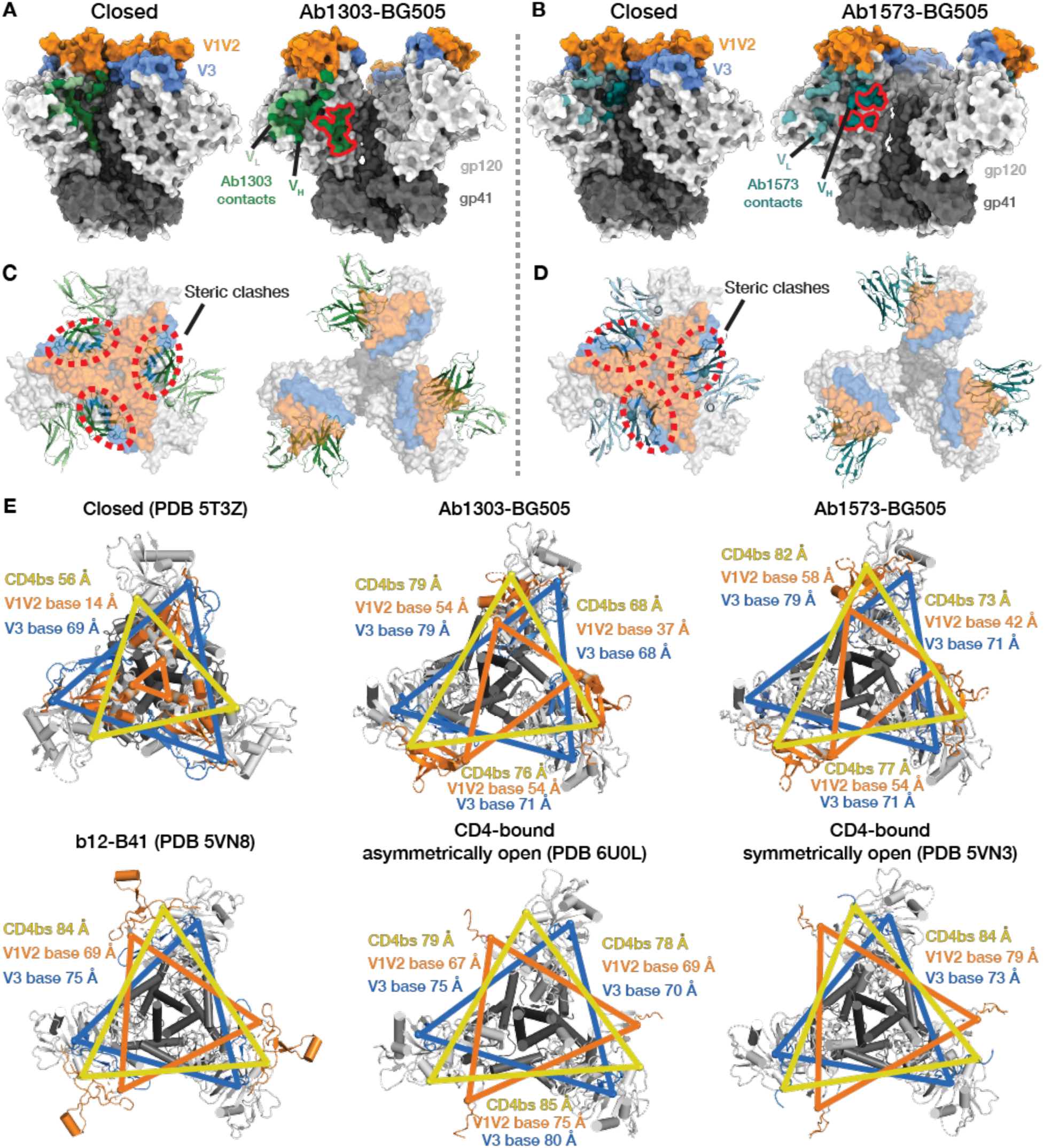
Ab1303 and Ab1573 bind an Env trimeric state distinct from the closed and CD4-bound open trimer conformations. (A,B) Ab1303 (A) or Ab1573 (B) contacts mapped onto a side-view surface representation of a closed BG505 trimer (PDB code 5CEZ) (left) and the occluded-open trimer to which each mAb binds (right). Regions of the antibody epitope that are buried in the closed state but accessible in the Ab1303- or Ab1573-bound state are outlined in red on the occluded-open Env trimer structures. (C,D) Cartoon representations of Ab1303 (C) or Ab1573 (D) interacting with closed (left) or occluded-open (right) Env trimers (seen from the top). Regions with steric clashes between the Fab and the closed trimer indicated by dashed, red ovals. (E) Comparison of inter-protomer distances of Cα atoms from selected residues at the CD4bs (yellow), the V1V2 base (orange), and the V3 base (blue) in a closed Env trimer (PDB 5T3Z), Ab1303-bound BG505 trimer, Ab1573-bound BG505 trimer, b12-bound B41 trimer (PDB 5VN8), a CD4-bound asymmetrically-open BG505 trimer (PDB 6U0L), and a sCD4-bound symmetrically-open B41 trimer (PDB 5VN3).

To compare and quantify outward displacements of gp120 protomers in different Env states, we measured inter-protomer distances of selected residues located in the CD4bs, V1V2 base, and V3 base of a closed BG505 Env trimer with analogous residues in BG505 trimers bound to Ab1303 or Ab1573 (Figure 5E). Inter-protomer distances were increased in the Ab1303- and Ab1573-bound trimers compared with the closed trimer, providing a quantitative measurement of openness. In addition, differences in the three inter-protomer distances for each measurement within the Ab1303- and Ab1573-bound Envs demonstrated trimer asymmetry compared with the symmetric closed trimer conformation. Comparisons with a structure of B41 SOSIP bound to b12 Fab^11^ showed that the Ab1303- and Ab1573-bound trimers resembled the b12-bound Env state more than the closed state, although the b12-bound trimer structure was three-fold symmetric (Figure 5E). Finally, measurements for all four of these trimers differed from the CD4-bound open state exemplified by the structure of a CD4-bound asymmetrically open BG505 trimer^14^ and a CD4-bound symmetrically open B41 trimer^11^ (Figure 5E).

Despite Env trimer opening, the gp120 V1V2 and V3 regions in the Ab1303-BG505 and Ab1573-BG505 structures exhibited only minor local structural rearrangements in which the gp120s were displaced as nearly rigid bodies from their central positions in the closed trimer structure (Figure 6A; Movie S1). Thus, most of each gp120 subunit, including the V1V2 and V3 regions, remained unchanged between the closed Env conformation and the open Ab1303- and Ab1573-bound Env trimers (Figure 6A, left and middle); the rmsd for superimposition of 347 gp120 Cα atoms (Gly41_gp120_ to Pro493_gp120_ excluding disordered residues) was ~1.3Å. By contrast, when bound to CD4, Env trimer opening did not result from rigid body rotations of gp120; instead the V1V2 loops were displaced from apex of each gp120 apex to the sides of the Env trimer to expose the coreceptor-binding site on V3, and portions of V1V2 and V3 were disordered^11,12,14^ (Figure 6A, right). In addition, the gp120 β2 and β3 strands at the beginning and end of the V1V2 loop switched positions with respect to their locations in closed, prefusion Env trimers to form a four-stranded anti-parallel β-sheet (4-stranded bridging sheet) (Figure 6B, right) instead of the 3-stranded sheet in closed Env structures (Figure 6B; left) (Movie S1). Notably, the ‘occluded open’ Env trimer conformations observed upon binding of Ab1303, Ab1573, or b12 included the 3-stranded sheet found in the closed, prefusion Env trimer rather than the 4-stranded bridging sheet in open, CD4-bound Env trimers (Figure 6B; middle; Movie S1).

**Figure 6.**
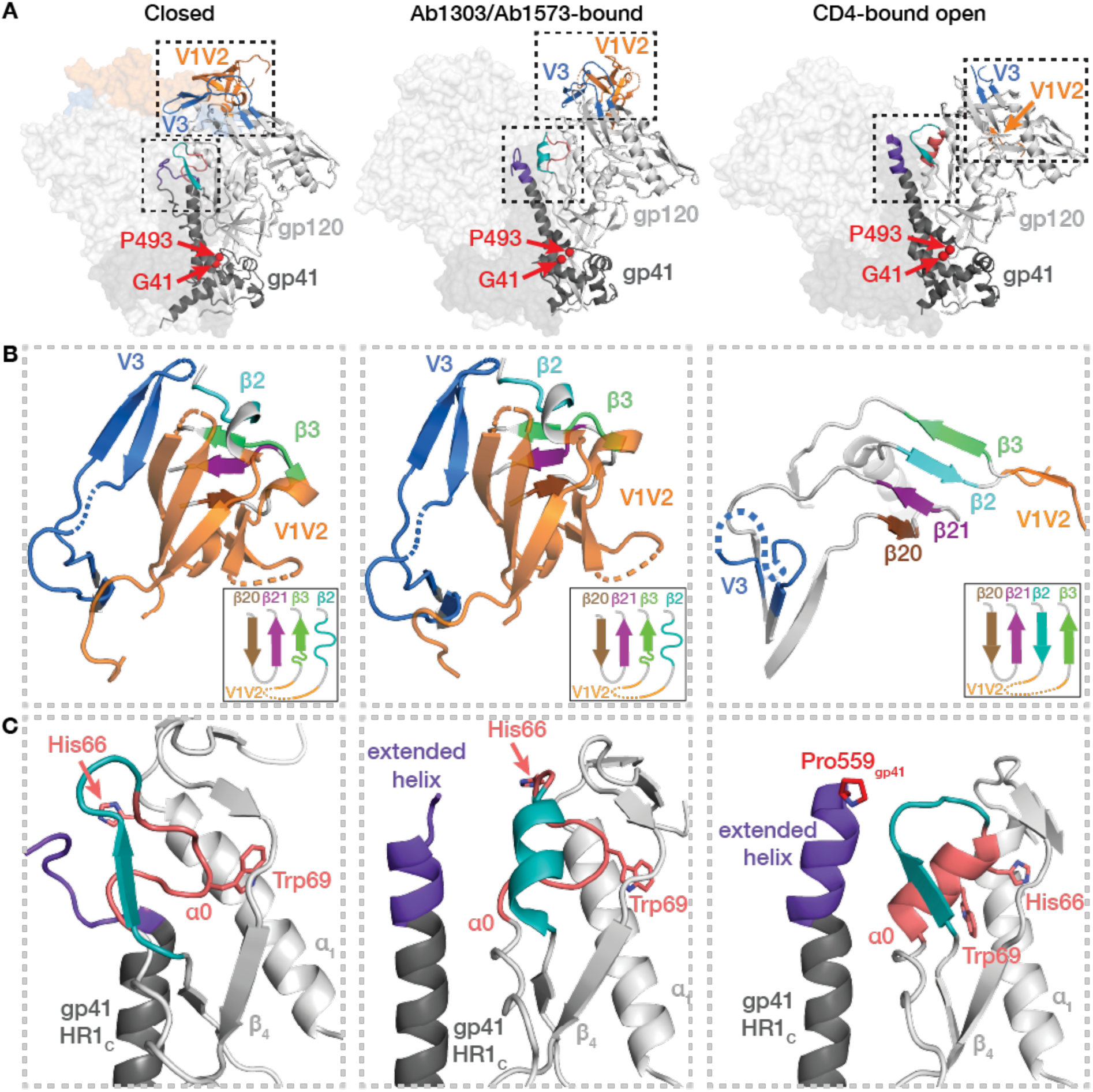
Structural changes between Env trimer states. (A) Conformations of gp120/gp41 protomers in closed (left), Ab1303/Ab1573-bound (middle), and CD4-bound (right) states. The gp120 V1V2-V3 regions and gp120 α_0_/gp41 HR1C regions are highlighted in dashed boxes. Positions of Gly41_gp120_ and Pro493_gp120_ are shown as red spheres. (B) Structural rearrangements of gp120 V1V2 and V3 regions in closed (left), Ab1303/Ab1573-bound (middle), and CD4-bound (right) states. Secondary structures and relative locations of the 3-stranded sheet (left and middle) and rearranged 4-stranded bridging sheet (right) are highlighted, and respective topology models are shown in insets. Dotted lines represent disordered regions. (C) Comparison of gp120 α_0_ and neighboring region in closed (left), Ab1303/Ab1573-bound (middle), and CD4-bound (right) structures. Residues Asp57_gp120_ - Glu62_gp120_ are teal, α_0_ residues (Lys65_gp120_ - Ala73_gp120_) are salmon, with the His66_gp120_ and Trp69_gp120_ side chains shown. The N-terminal segment of gp41 HR1_C_ is colored dark purple.

Local structural rearrangements in Asp57_gp120_ - Ala73_gp120_, residues that are immediately N-terminal to the gp120 α_0_ region (residues Lys65_gp120_ - Ala73_gp120_), provided further evidence that the Ab1303- and Ab1573-bound trimers adopted a state distinct from the closed state recognized by CD4bs bNAbs: in the closed state, residues Asp57_gp120_ - Glu62_gp120_ formed a β-strand and a short loop (Figure 6C, left), whereas they formed a two-turn α-helix and the α_0_ residues remained as a loop in Ab1303/Ab1573-bound open conformation (Figure 6C, middle). By contrast, in the CD4-bound fully-open state, the Asp57_gp120_ - Glu62_gp120_ segment formed a β-strand and short loop; whereas the α_0_ segment (residues Lys65_gp120_ - Ala73_gp120_), which was a loop in both the closed and the Ab1303/Ab1573-bound open states, was an α-helix in the CD4-bound fully-open state (Figure 6C, right). The structural rearrangement of α_0_ was accompanied by protein sidechain repositioning: in the closed and Ab1303/1573-bound open states, the Trp69_gp120_ sidechain was sandwiched between the α1 helix and β_4_ strand and His66_gp120_ was solvent exposed, whereas in the CD4-bound open state, the Trp69_gp120_ sidechain was rearranged such that the His66_gp120_ sidechain occupied a nearly analogous position (Figure 6C).

The conformation of the C-terminal portion of gp41 heptad repeat segment 1 (HR1C) also exhibited changes between the closed, Ab1303- and Ab1573-bound open, and CD4-bound open Env trimer states. In closed prefusion Env trimers, residues N-terminal to Thr569_gp41_ adopted a loop structure (Figure 6C, left). Outward rotations of the gp120 subunits in the Ab1303-/Ab1573-bound open Env created space for the gp41 HR1_C_ to extend its three-helix coiled-coil structure, lengthening the α-helices by 1.5 turns (Figure 6C, middle). Additional outward gp120 rotations combined with V1V2 and V3 displacements created more space for the central α-helices in the CD4-bound fully-open state; thus gp41 residues that were disordered in the closed and occluded open trimer states extended the HR1C N-terminal helical structure by another helical turn, with ordered residues terminating at around residue Pro559_gp41_^11–14^, the site of the Ile-to-Pro stabilizing mutation in SOSIPs^9^ (Figure 6C, right). Thus, the Ab1303- and Ab1573-bound Env structures revealed an occluded-open trimer state distinct from both the closed, prefusion and the CD4-bound fully-open trimer conformations.

### DEER suggests that the unliganded Env trimers contain both occluded-open and other Env conformations

To evaluate the conformational flexibility of ligand-free and antibody-bound trimer in solution, we used double electron-electron resonance (DEER) spectroscopy to probe inter-protomer distances between V1V2 regions in different Env trimer states. DEER can be used to derive distances between electron spin pairs ranging from 17-80Å by detecting their respective dipolar interactions^36^. By recording a snapshot of the equilibrium distance distributions of flash-frozen samples, DEER data report molecular motions in solution to provide insight into conformational heterogeneity. We previously used DEER to evaluate spin-labeled BG505 and B41 SOSIPs in the presence and absence of antibodies, CD4, and a small molecule ligand, finding a relatively homogeneous trimer apex, more conformational heterogeneity at the trimer base, and inter-protomer distances between spin labels that were consistent with bNAb-bound closed Env structures and CD4-bound open Env structures^37^.

In the present studies, we introduced a free cysteine into a gp120-gp41 protomer of the BG505 SOSIP in order to use site-directed spin labeling^38^ to covalently attach a nitroxide spin label with a V1 side chain^39^. This approach results in three spin labels on each Env, which form a triangle of spin labels, either equilateral, isosceles, or scalene depending on whether the labeled Env adopts a three-fold symmetric or asymmetric conformation. Thus, DEER measurements in a conformationally-rigid Env trimer would report one distance in a symmetric Env and two or more distances in asymmetric Envs. The most probable distance in a DEER distribution is defined by the largest peak area and represents the dominant structural state in a population of states. The presence of multiple peaks in a DEER distribution indicates conformational heterogeneity, with individual peak widths related to the flexibility of that conformation and of the attached spin label^38,40^. In general, peaks representing 17–65Å distances can be assigned with confidence, whereas distances > 65Å are detected with less accuracy^36^.

To choose a site for spin labeling, we used the Ab1303- and Ab1573-Env structures to identify solvent-exposed residues, which when spin-labeled, would result in distinguishable inter-protomer distances in different Env conformations. We also restricted candidate sites to residues located in a β-sheet to minimize potential flexibility of the attached spin label and excluded residues that were involved in interactions with other residues to avoid disrupting protein folding. The optimal candidate residue, V1V2 residue Ser174_gp120_, fulfilled these criteria, with inter-protomer Cα-Cα distances measured as 38Å in a closed Env structure, ranging from 40Å-60Å in the asymmetric Ab1303- and Ab1573-bound Env structures, 67Å in a b12-bound Env, and ~157Å (far out of DEER range) in a CD4-bound open Env (Figure 7A). Although the V1 spin label is small (about the size of an amino acid) and contributes limited width to DEER distance distributions^41^, distances between spin label side chains measured by DEER only rarely equal the Cα-Cα inter-protomer distance since the radical center is found on the nitroxide ring, not the peptide linkage. As such, DEER results can be complicated by conformational heterogeneity and flexibility intrinsic to the protein studied.

**Figure 7.**
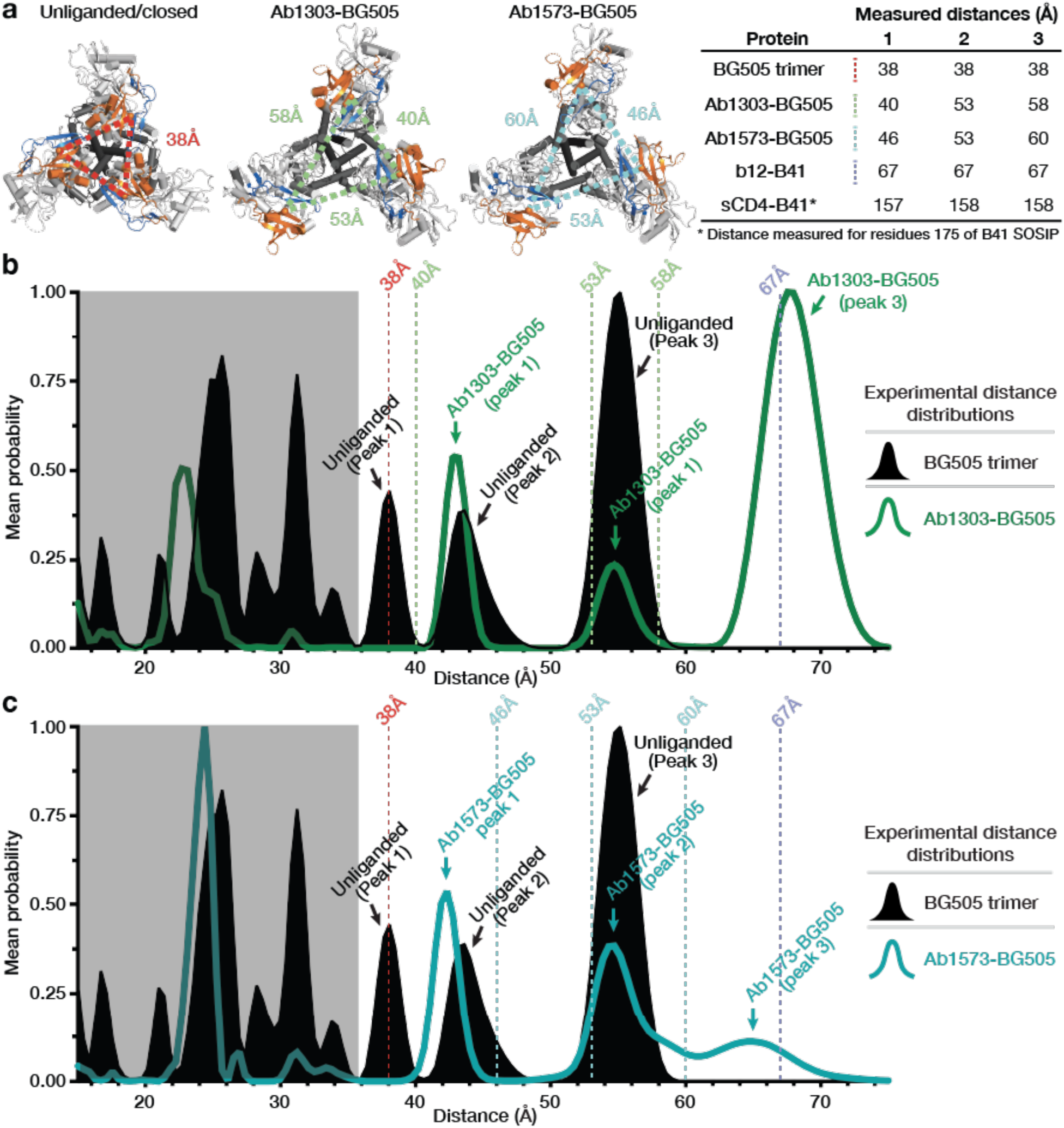
DEER spectra of unliganded BG505 trimer compared to Ab1303-bound and Ab1573-bound BG505 trimer. (A) Left: Triangles showing distances between residue 174 Cα atoms on cartoon representations of Env trimers seen from the top. Right: Inter-protomer distances between Cα atoms for V1V2 residue 174 in different Env conformational states measured using coordinates from the indicated structures (BG505 trimer: PDB 5T3Z; Ab1303-BG505 and Ab-1573-BG505: this study; b12-B41: PDB 5VN8; CD4-B41: PDB 5VN3). (B,C) Left: Distance distributions for spin labels attached to V1V2 residue 174 in unliganded BG505 Env (solid black), Ab1303-bound BG505 (green trace in B), and Ab1573-bound BG505 (teal trace in C). Vertical dashed lines indicate the inter-subunit distances measured from coordinates (panel A) for each site in structures of closed, unliganded Env (red line), Ab1303-bound Env (green dashed lines in B), Ab-1573-bound Env (teal dashed lines in C), and b12-bound Env (purple dashed line). Right: Legend for DEER spectra.

The S174C mutant version of BG505 SOSIP was expressed, purified, and labeled with the V1 side chain. Ab1303 and Ab1573 Fabs were added at a 3 molar excess to V1-labeled BG505, and liganded and unliganded samples were incubated at 37℃ for three hours before being flash-frozen in liquid nitrogen for resonance measurements. The DEER spectrum of unliganded BG505 (black trace in Figure 7B,C) showed a complicated collection of peaks, indicating conformational heterogeneity of the V1V2 region in the vicinity of gp120 residue 174. These results differ from previous DEER experiments from which we derived BG505 V1V2 distances from spectra recorded after incubation at 4℃, which showed a more homogeneous distance distribution with a dominant peak observed at the expected inter-protomer distance^37^. In the present experiments, one of the major peaks, centered at ~38Å, corresponds to the residue 174 inter-protomer distance in a closed BG505 Env structure (Figure 7B,C; red vertical line). The other major peaks for the unliganded BG505 sample, including major peaks at distances between ~20Å and ~35Å, were not readily interpretable based on closed SOSIP Env structures. However, the presence of the inter-protomer distances other than 38Å suggests that the unliganded BG505 SOSIP trimer can adopt conformational states in addition to the known closed, prefusion conformation. The broad heterogeneity of conformational states seen here may have been induced by incubation at 37℃.

We also collected DEER data for BG505 complexes with Ab1303 and Ab1573 (green and cyan traces in Figures 7B and C, respectively). Some of the short distance peaks, most notably a single major peak at ~24Å, were observed for both antibody-bound Envs (Figure 7B,C). In addition to this structurally uninterpretable peak, also present in both spectra were peaks at ~43Å and ~55Å, likely corresponding to the structurally-measured inter-protomer distance of 40Å/46Å (measured distance 1 for Ab1303-Env and Ab1573 Env complexes) and a combination of the 53Å/58Å (Ab1303) and 53Å/60Å (Ab1573) distances (measured distances 2 and 3; green vertical lines in Figure 7B; cyan vertical lines in Figure 7C). Peaks at or close to these distances were found in the unliganded BG505 DEER spectrum, suggesting that the conformational states observed for Ab1303/Ab1573-binding Env also exist at a lower population in unliganded BG505. Interestingly, peaks near 67Å – the measured inter-protomer distance for residue 174 in a b12-bound Env trimer – are observed in both the Ab1303-bound and Ab1573-bound Envs (major peak in Ab1303-bound Env spectrum; a minor peak in the Ab1573-bound spectrum), suggesting that binding of these antibodies induced a sub-population of Envs with a b12-bound conformation that was not captured in the cryo-EM structures.

## Discussion

Many HIV-1 vaccine efforts focus on using soluble Env trimers as immunogens to raise bNAbs^42^. Here we characterize Ab1303 and Ab1573, two CD4bs bNAbs raised by sequential immunization in NHPs of SOSIP trimer immunogens attached to VLPs^19^. Unexpectedly, both antibodies bind the CD4bs of an open form of HIV-1 Env trimer rather than the closed, prefusion state typically targeted by bNAbs raised in humans by natural infection^10^. In common with CD4-bound Envs, the trimer conformation recognized by Ab1303 and Ab1573 includes outward rotations of gp120, but the gp120 V1V2 loops are not rearranged to expose the coreceptor binding site on V3; thus, the Env trimer is open but the co-receptor binding site is occluded. The Ab1303/Ab1573-bound occluded-open Env trimer conformation shares structural features with the conformation recognized by b12^11^, an early CD4bs bNAb isolated from a phage display library^15^. Of relevance to immunogen design efforts is whether the occluded-open Env conformation exposes new epitopes that might elicit off-target non-neutralizing antibodies against trimer surfaces that would be buried in closed, prefusion Env trimers. Comparisons of BSAs between closed and occluded-open Env structures show that regions of V1V2 that are inaccessible in closed trimers (purple in Figure S3) might be accessible for antibody binding in the occluded-open Env conformation.

The question of which features of an Env-binding ligand induce an open coreceptor-binding HIV-1 Env conformation is prompted by the existence of two distinct open trimer conformations (Figure 6A): (*i*) the CD4-bound open trimer, a coreceptor-binding conformation in which V1V2 relocates to the sides of the Env trimer to expose V3 and form a 4-stranded gp120 bridging β-sheet^11–14^, versus (*ii*) the occluded-open trimer in which the gp120s rotate outwards, but V1V2 remains “on top” of gp120 to shield the coreceptor binding site. Structures of the b12-Env^11^ and the Ab1303/Ab1573-Env complexes reported here demonstrate that Env opening through gp120 rotation is not sufficient to induce the further structural rearrangements associated with CD4 binding (Figure 6B,C). One difference that distinguishes CD4 from b12, Ab1303, Ab1573 and most other CD4bs bNAbs is that the antibodies lack a counterpart of CD4 residue Phe43, which inserts into the “Phe43” pocket on gp120^43^. We showed that small molecule CD4 mimetic entry inhibitors that insert into the gp120 Phe43 pocket recognize the CD4-bound open trimer conformation^35^, whereas CD4 mimetics drugs that bind orthogonally to the Phe43 pocket bind closed, prefusion Envs^44^. Interaction with the gp120 Phe43 pocket may be necessary for recognition of the CD4-bound open trimer conformation, but is unlikely to be sufficient since CD4bs bNAbs such as N6^45^ contain a CD4 Phe43 counterpart within their CDRH2 region, yet bind closed, prefusion Env trimers^46^.

Our findings suggest that portions of the Ab1303 and Ab1573 epitopes on gp120 are buried on a closed, prefusion Env trimer (Figure 5A) and that there are potential steric clashes between the Fabs and Env when they are docked onto their respective binding sites of closed Env (Figure 5C,D). This implies that the Env trimer conformation that triggers development of this type of CD4bs bNAb is similar or equivalent to the occluded-open Env conformation described here. Indeed, DEER spectroscopy experiments suggested that a population of unliganded BG505 SOSIP Envs that had been incubated at 37°C adopted a conformation consistent with the occluded-open conformation recognized by Ab1303 and Ab1573 (Figure 7), therefore this conformation may have been present on at least a subset of the SOSIP-based immunogens used in the sequentially-immunized NHPs from which these antibodies were derived^19^. In addition, the Env trimers of HIV-1 strains that are neutralized by Ab1303 and Ab1573^19^ may more readily adopt the occluded-open conformation than Envs in neutralization-resistant strains.

The DEER results, together with the demonstration of temperature-dependent changes in Ab1303 and Ab1573 binding stoichiometry, suggest that physiological temperature facilitates conformational changes in soluble Env trimers that result in the occluded-open state. Our previous DEER studies to probe conformations of Env SOSIPs conducted with 4°C incubations concluded that unliganded SOSIPs showed conformations that were consistent with the closed pre-fusion trimer conformation, with no evidence for the CD4- or b12-bound open states^37^. For example, DEER spectra of the unliganded BG505 SOSIP labeled in V1V2 (residue 173) prepared at 4℃ showed a dominant inter-protomer distance signal between 30-40Å, consistent with distances measured for the closed Env conformation^37^. In this study, the comparable unliganded BG505 SOSIP sample labeled in V1V2 (residue 174) prepared at 37℃ reported interspin distances consistent with the Ab1303/Ab1573-bound occluded-open trimer conformation. This suggests that at 37℃, the temperature at which antibodies are generated *in vivo*, Env trimers attached to VLPs can adopt different conformational states between defined closed and open conformations. Whether the occluded-open conformation is present on membrane-bound viral Env trimers remains unknown, but the isolation of the b12 bNAb from a phage display library constructed using bone marrow from an HIV-1–infected individual is consistent with the idea that viruses include Envs with this or a similar conformation. The discovery that the b12-bound conformation of HIV-1 Env trimer is recognized by vaccination-induced neutralizing antibodies suggests the occluded-open conformation of HIV-1 as a potential target for immunogen design.

## Data Availability

The atomic models have been deposited in the Protein Data Bank (PDB) accession codes 7RYU for Ab1303 Fab and 7RYV for Ab1573 Fab, xxxx for Ab1303-BG505 and xxxx for Ab1573-BG505 complex. The cryo-EM maps have been deposited in the Electron Microscopy Data Bank (EMDB) with entries xxxx and xxxx for Ab1303-BG505 and Ab1573-BG505 complexes, respectively.

## Acknowledgements

We thank Anthony P. West (Caltech) for help with analysis of antibody sequences. Cryo-EM was performed in the Beckman Institute Resource Center for Transmission Electron Microscopy at Caltech with assistance from directors A. Malyutin and S. Chen. We thank Jost Vielmetter and Pauline Hoffman at the Beckman Institute Protein Expression Center at Caltech for protein production, John Moore (Weill Cornell Medical College) for the BG505 stable cell line, Kristie M. Gordon (The Rockefeller University) for assistance with flow cytometry, and Rogier W. Sanders and Marit J. van Gils (Academisch Medisch Centrum Universiteit van Amsterdam) for providing AviTagged and biotinylated BG505 and B41 SOSIP trimers for sorting. This work was supported by the National Institute of Allergy and Infectious Diseases (NIAID) HIVRAD P01 AI100148 (to P.J.B. and M.C.N.), Gates CAVD grant INV-002143 (to P.J.B., M.A.M., and M.C.N.), the Intramural Research Program of the National Institute of Allergy and Infectious Diseases, NIH. (R.G. and M.A.M), NIH P50 AI150464 (P.J.B.). M.C.N. is an HHMI Investigator.

This is a preprint of an article published in *Nature Communications*. The peer-reviewed, authenticated version is available online at: https://doi.org/10.1038/s41467-022-28424-3.

## Author contributions

Z.Y., K.A.D., M.A.M., M.C.N., W.L.H., and P.J.B. designed the research. Z.Y., K.A.D., M.D.B., M.A.G.H., H.B.G., A.T.D, A.E., and R.G. performed experiments and analyzed results. Z.Y., K.A.D., M.D.B., and P.J.B. wrote the manuscript with input from co-authors.

## Methods

### Protein expression and purification

The native-like, soluble HIV Env gp140 trimer BG505 SOSIP.664 construct with ‘SOS’ mutations (A501C_gp120_, T605C_gp41_), the ‘IP’ mutation (I559P_gp41_), *N*-linked glycosylation site mutation (T332N_gp120_), an enhanced furin protease cleavage site (REKR to RRRRRR), and truncation after the C-terminus of gp41 residue 664^9^ was cloned into pTT5 vector (National Research Council of Canada) and expressed in transiently-transfected Expi293F cells. BG505 Env trimer was purified from transfected cell supernatants as described^47^ by 2G12 immunoaffinity and size-exclusion chromatography (SEC) with a Superose 6 16/600 column followed by Superdex 200 Increase 10/300 GL column (GE Life Sciences).

The heavy and light chains of 6x-His tagged Ab1303 and Ab1573 Fabs were expressed in transiently-transfected Expi293F cells and purified by Ni-NTA chromatography followed by SEC as described^19^. IgG proteins were expressed in transiently-transfected Expi293F cells and purified by protein A affinity chromatography (GE Healthcare) followed by SEC as described^21,47^.

### Competition ELISA

BG505 SOSIP trimers were randomly biotinylated using the EZ-Link NHS-PEG4-Biotin kit (Thermo Fisher Scientific) according to the manufacturer’s guidelines. Based on the Pierce Biotin Quantitation kit (Thermo Fisher Scientific), the number of biotin molecules per protomer was estimated to be 1.5. Biotinylated BG505 SOSIP timers were immobilized on Streptavidin-coated 96-well plates (Thermo Fisher Scientific) at a concentration of 5 μg/mL in blocking buffer (1% BSA +1% goat serum in TBS-T) for 1 hour at room temperature (RT). After washing plates once in TBS-T, plates were incubated with a Fab derived from a bNAb that targets the V3 loop (10-1074), the fusion peptide (VRC34), the V1V2 loop (PG16), or the CD4bs (3BNC117), at a concentration of 100 μg/mL in blocking buffer for 1 hour at RT. After washing plates twice in TBS-T, a concentration series of Ab1303 or Ab1573 IgG was added to the Fab-BG505 complexes with a top concentration of 100 μg/mL in blocking buffer and 4-fold dilutions for 1 hour at 37°C. After washing plates three times in TBS-T, bound IgG was detected using an HRP-conjugated goat anti-human Fc antibody (Southern Biotech) at a dilution of 1:10,000 in blocking buffer for 1 hour at RT. After washing plates three times with TBS-T, 1-Step Ultra TMB substrate (Thermo Fisher Scientific) was added for 5 min and plates were analyzed using a plate reader (BioTek).

### PGT145 capture ELISA

This capture ELISA was performed as described previously with minor modifications^9^. Briefly, Corning Costar 96-Well Assay high binding plates (07-200-39) were coated and incubated overnight at 4°C with PGT145 IgG at 5 μg/ml in 0.1 M NaHCO_3_ (pH 9.6). Unbound PGT145 IgG was removed, and wells were blocked with 3% BSA in TBS-T (20mM Tris, 150 mM NaCl, 0.1% Tween20) for 1 hour at room temperature. BG505 SOSIP.664 was added at 10 μg/ml and incubated for 1 hour at room temperature, then removed. Serially diluted Fabs in 3% bovine serum albumin in TBS-T were added, incubated at room temperature for 2 hours, then washed three times with TBS-T. Horseradish peroxidase labeled mouse anti-His tag antibody (GenScript: A00186) was added for 30 minutes at 1:1000 dilution, followed by 3 washes with TBS-T. 1-Step™ Ultra TMB-ELISA Substrate Solution (ThermoFisher Scientific: 34029) was added for colorimetric detection. Color development was quenched with 1.0 N HCl, and absorption was measure at 450 nm. Two independent, biological replicates were performed.

### X-ray crystallography

Crystallization screens for Ab1303 Fab and Ab1573 Fab were performed using the sitting drop vapor diffusion method at room temperature by mixing concentrated Fabs with an equal amount of reservoir solution (Hampton Research) using a TTP Labtech Mosquito automatic microliter pipetting robot. Ab1303 Fab crystals were obtained in 10% (v/v) PEG 200, 0.1M Bis-Tris propane (pH 9.0), and 18% w/v PEG 8,000. Ab1573 Fab crystals were obtained in 0.1M Tris (pH 8.2), 26% (w/v) PEG 4000. Ab1303 Fab crystals were directly looped and cryopreserved in liquid nitrogen, whereas Ab1573 Fab was briefly mixed with 15% glycerol cryoprotectant solution before cryopreservation in liquid nitrogen.

X-ray diffraction data were collected at Stanford Synchrotron Radiation Lightsource (SSRL) beamline 12-1 equipped with an Eiger X 16M pixel detector (Dectris) at a wavelength of 0.97946Å. Recorded data were indexed, integrated, scaled in XDS^48,49^ and merged with AIMLESS v0.7.4^50^. The structure of Ab1303 Fab was determined by molecular replacement using PHASER v2.8.2^51^ and a search model comprising separate V_H_-V_L_ and C_H_1-C_L_ domains of a human antibody (PDB 4YK4) with the CDR loops removed. The structure of Ab1573 Fab was determined similarly, except using the Ab1303 Fab as the search model. Coordinates of both Fabs were refined using Phenix^52,53^ and iterations of manual building in Coot^54^ (Supplementary Table 2).

### Cryo-EM sample preparation

Ab1303-BG505 and Ab1573-BG505 complexes were prepared by incubating purified and concentrated Ab1303 and Ab1573 Fabs with BG505 SOSIP.664 trimer at a molar ratio of (3.6:1 Fab:Env) at 37°C for 2 hours. A final concentration of 0.05% (w/v) florinated octylmaltoside (Anatrace) was added to both samples immediately before cryo-freezing. Cryo-EM grids were prepared using a Mark IV Vitrobot (ThermoFisher) operated at 12°C and 95% humidity. 2.6 μL of concentrated sample was applied to 300 mesh Quantifoil R1.2/1.3 grids, blotted for 4 seconds, grids were then vitrified in liquid ethane.

### Cryo-EM data collection and processing

Cryo-grids were loaded onto a 200kV Talos Arctica electron microscope (ThermoFisher) (Ab1303-BG505 complex) or a 300kV Titan Krios electron microscope (ThermoFisher) equipped with a GIF Quantum energy filter (slit width 20 eV) operating at a nominal 105,000x magnification (Ab1573-BG505). For data collection on the Krios, defocus ranges for both Ab1303-BG505 and Ab1573-BG505 datasets were set to 1.4-3.0 μm. Movies were recorded using a 6k x 4k Gatan K3 direct electron detector operating in super-resolution mode with a pixel size of 0.869 ÅApixel^−1^ (Arctica) or 0.855 Å•pixel^−1^ (Krios) and collected using SerialEM v3.7 software. The recorded movies were sectioned into 40 subframes with a total dose of 60 e^−^•Å^−2^, generating a dose rate of 1.5 e^−^/Å^2^•subframe. A total of 8,036 (Ab1303-BG505) and 8,478 (Ab1573-BG505) movies were motion-corrected using MotionCor2^55^ with 2x binning. The CTFs of motion-corrected micrographs were estimated using CTFFIND v4.1.14^56^. For both datasets, a set of ~1000 particles were manually picked and reference-free 2D classes were selected for automatic particle picking using RELION AutoPicking^57,58^. Automatically-picked particles were subjected to iterations of reference-free 2D class averaging. A closed-conformation trimer (PDB 5CEZ) map that was low-pass filtered to 80Å was used as reference for 3D classifications and high-resolution 3D refinement in RELION v3.1^57,58^. CTF refinements were performed on particles used previously in 3D refinement, and the CTF-refined particles were subsequently polished and subjected to a last iteration of 3D refinement and map sharpening. 3D FSCs of maps were calculated using the Remote 3DFSC Processing Server as described^59^. Local resolutions of the refined maps were calculated using RELION v3.1^57,58^.

### Model building

Coordinates for gp120, gp41, Ab1303 Fab, and Ab1573 Fab were fitted into the corresponding regions of the density maps. The following coordinate files were used for initial fitting: BG505 gp120 monomer (PDB 5T3Z), gp41 monomer (PDB 5T3Z), and crystal structures of Ab1303 and Ab1573 (this study). Coordinates for the two Fab-BG505 structures and *N*-linked glycans were manually refined and built in Coot^54^. Iterations of whole-complex refinements using *phenix.real_space_refine*^52,53^ and manual refinements were performed to correct for interatomic bonds and angles, clashes, residue side chain rotamers, and residue Ramachandran outliers.

### Structural analyses

CDR lengths were derived based on IMGT definitions^60^. Structural figures were made using PyMOL (Schrödinger, LLC) or ChimeraX^61^. Buried surface areas were calculated using PDBePISA^62^ and a 1.4 Å probe. Potential hydrogen bonds were assigned using the geometry criteria with separation distance of <3.5 Å and A-D-H angle of >90 ̊. The maximum distance allowed for a potential van der Waals interaction was 4.0 Å. Protein surface electrostatic potentials were calculated in PyMOL (Schrödinger LLC). Briefly, hydrogens were added to proteins using PDB2PQR^63^, and an electrostatic potential map was calculated using APBS^64^. Epitopes for antibodies in Figure 3 were identified as gp120 residues containing an atom within 4 Å of an antibody as calculated in PyMOL (Schrödinger, LLC).

### SOSIP V1 Spin Labeling and Pulsed DEER Spectroscopy

SOSIP spin labeling and pulsed DEER spectroscopy were performed similarly to methods described previously^37^. Briefly, purified SOSIP proteins where concentrated to ~100 μM in TBS (pH 7.4) and reduced with tris(2-carboxyethyl)phosphine (TCEP) buffer such that the final concentration of TCEP was in a 2x molar excess relative to the target cysteine residue for one hour. TCEP was removed using a desalting column (Zeba, 89883) and the reduced protein was then incubated with 5 molar excess of the V1 nitroxide spin label (bis(2,2,5,5-tetramethyl-3-imidazoline-1-oxyl-4-il)-disulfide) for 5 hours at room temperature then at 4°C overnight. A size exclusion chromatography column (Superose 6 10 300) was used to remove excess V1 spin label. V1-labeled SOSIP was then buffer exchanged into deuterated solvent containing 20% glycerol. Unliganded V1-labeled SOSIP and V1-labeled SOSIP incubated with a 3 molar excess of Ab1303 Fab or Ab1573 Fab were placed at 37°C for 3 hours immediately prior to flash freezing.

For DEER spectroscopy, approximately 60 μL samples of *~*150 μM spin-labeled protein complexes were flash frozen within a 2.0/2.4 mm borosilicate capillary (Vitrocom, Mountain Lakes, NJ) in liquid nitrogen. Sample temperature was maintained at 50 K during data collection by a recirculating/closed-loop helium cryocooler and compressor system (Cold Edge Technologies, Allentown, PA). Four-pulse DEER spectroscopy data were collected on a Q-band Bruker ELEXSYS 580 spectrometer using a 150 W TWT amplifier (Applied Engineering Systems, Fort Worth, TX) and an E5106400 cavity resonator (Bruker Biospin). Pulse lengths were optimized via nutation experiment but ranged from 21 to 22 ns (π/2) and 42 to 44 ns (π); Observer frequency was set to a spectral position 2 G downfield of the low and central resonance intersection minimum in the absorption spectrum, and the pump envelope frequency was a 50 MHz half-width square-chirp pulse (generated by a Bruker arbitrary waveform generator) set 70 MHz downfield from the observer frequency. Dipolar data were analyzed using LongDistances v.932, a custom program written by Christian Altenbach in LabVIEW (National Instruments); software available online (http://www.biochemistry.ucla.edu/biochem/Faculty/Hubbell/) and described elsewhere^65^.

---

**Figure S1.**
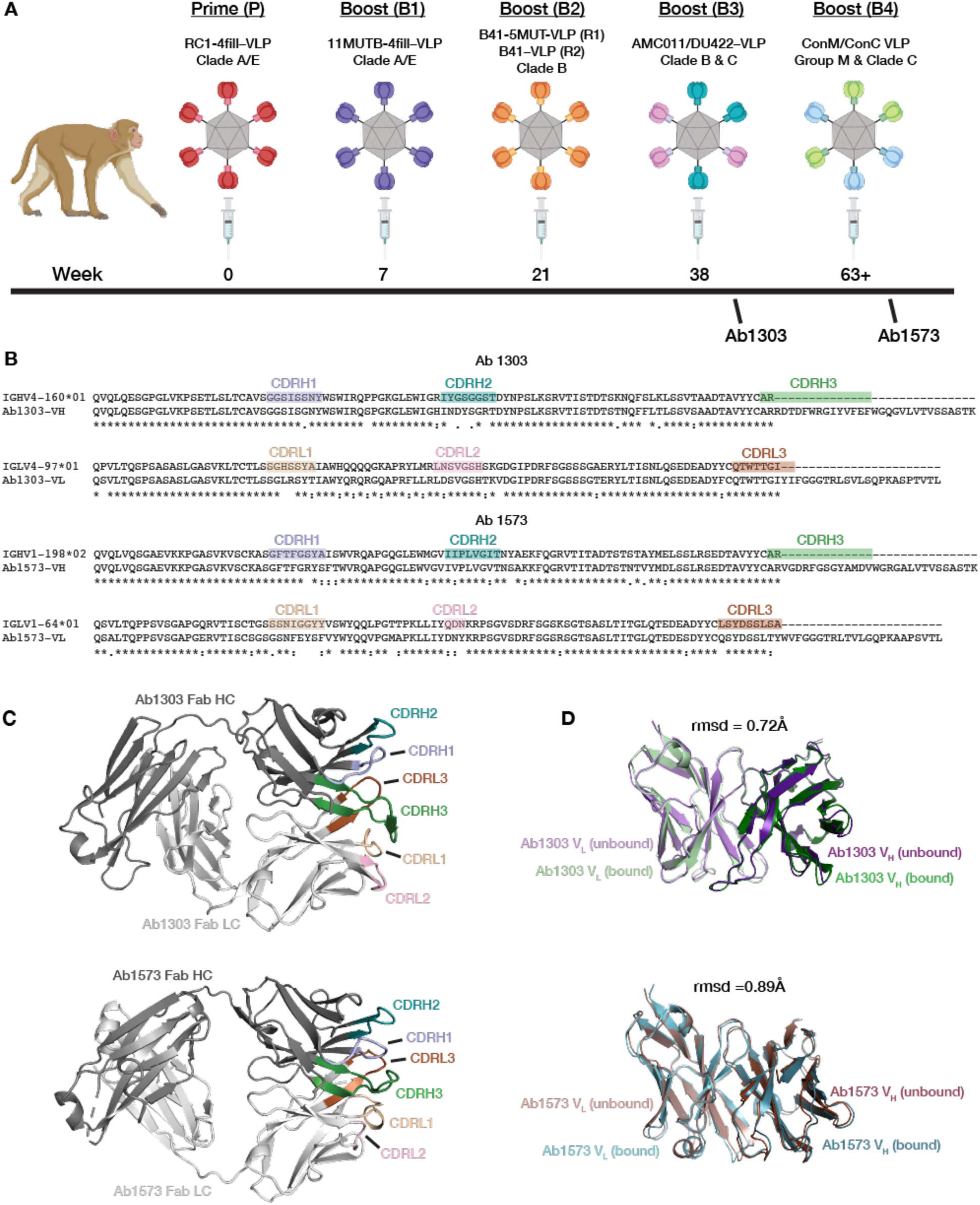
Characterization of Ab1303 and Ab1573. (A) Schematic of sequential immunization in NHPs from which Ab1303 (Regimen 1, R1) and Ab1573 (Regimen 2, R2) were derived. Ab1303 was isolated from NHP 4 (T15) after Boost 3; Ab1573 was isolated from NHP 1 (DGJI) after Boost 4 as described in^19^. (B) Sequence alignments of Ab1303 and Ab1573 V_H_ and V_L_ domains with their germline V gene precursors. (C) Crystal structures of unliganded Ab1303 and Ab1573 Fabs. CDRs of the two antibodies were highlighted in various colors. (D) Superimposition of structures of bound (from Fab-Env cryo-EM structures) and free (from Fab crystal structures) V_H_-V_L_ domains.

**Figure S2.**
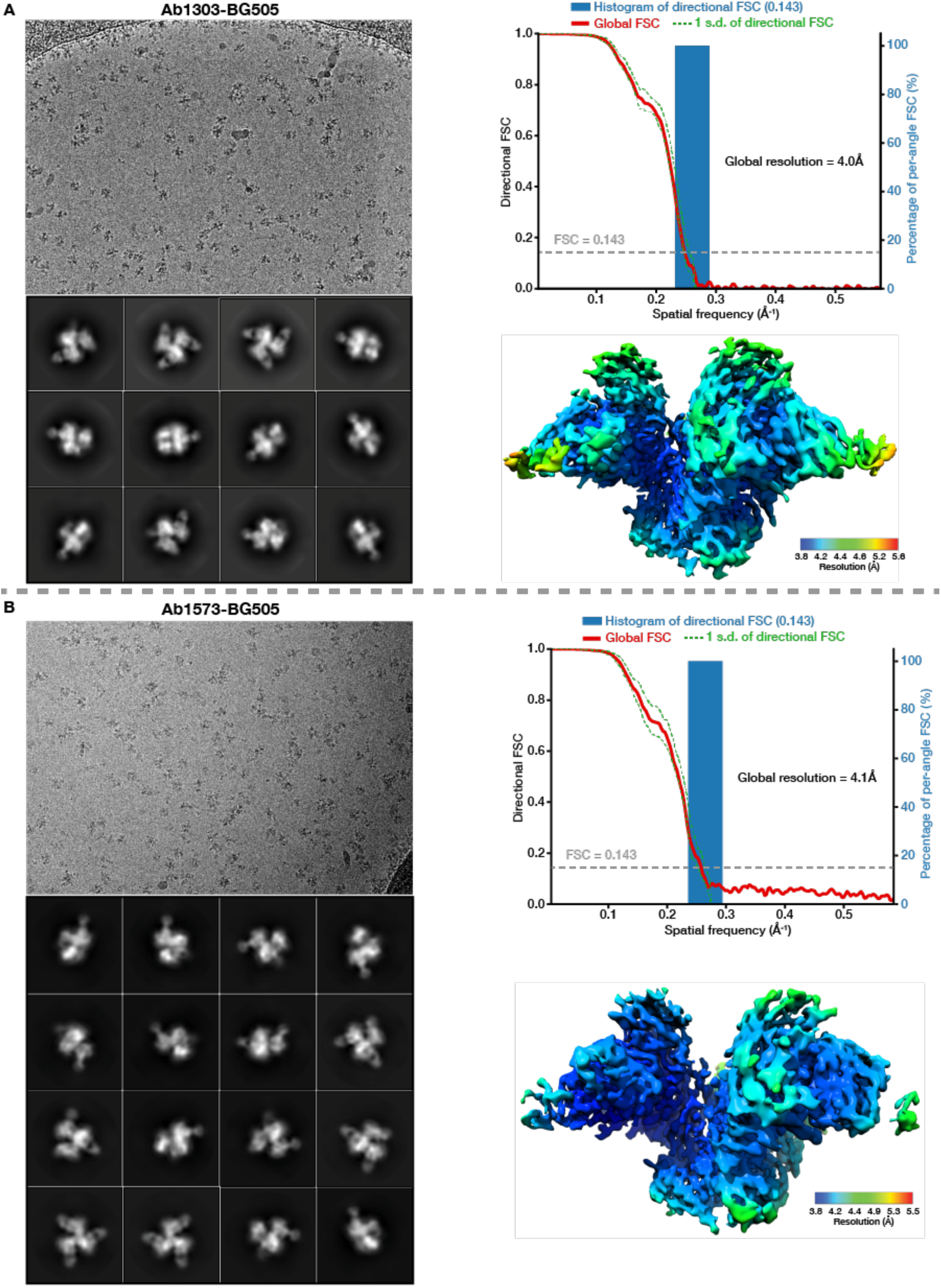
Cryo-EM processing, validation and reconstructions. (A,B) Left: Example micrographs and 2D class averages of Ab1303-BG505 (A) and Ab1573-BG505 (B) complexes. Upper right: Plots of global half-map FSCs (solid red line), directional resolution values ±1σ from the mean (left axis, green dashed lines), and histogram distributions sampled over 3D FSC (right axis, blue bars) for Ab1303-BG505 (B) and Ab1573-BG505 (B). Bottom right: local resolution maps of Ab1303-BG505 (A) and Ab1573-BG505 (B).

**Figure S3.**
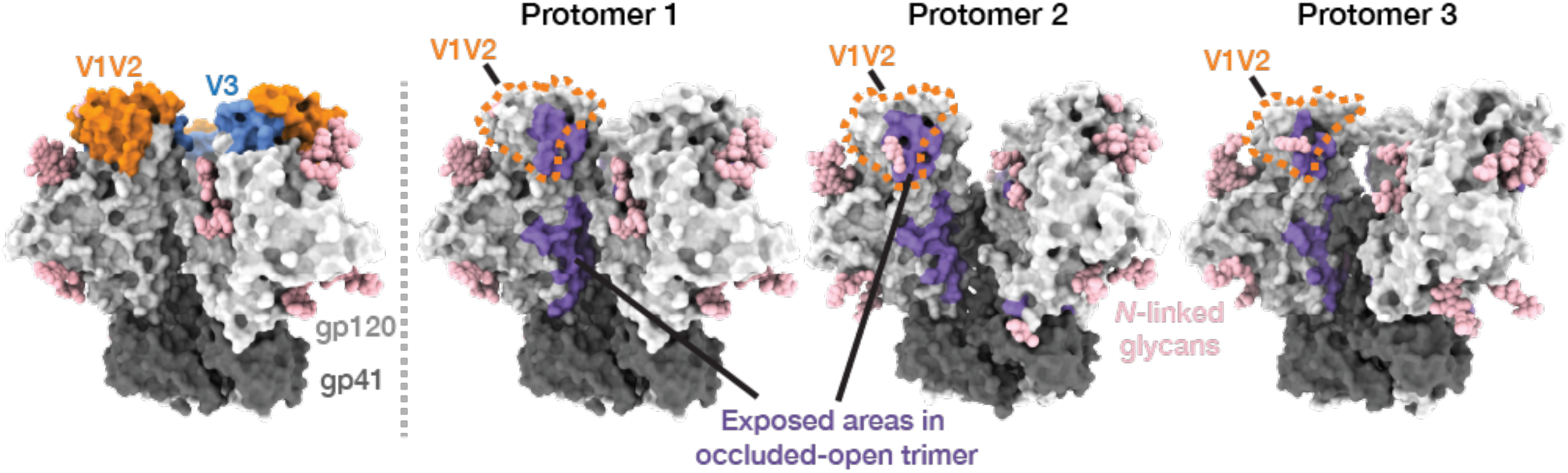
gp120 surface area exposed in occluded-open versus closed Env trimers. Left: Occluded-open Env trimer showing V1V2 and V3 regions as colored highlights. Right: Env trimers showing surface areas (purple) that are exposed in occluded-open trimers but buried in closed trimers. Orange dashed shape indicates V1V2 region.

**Supplementary table 1.** X-ray data collection and refinement statistics (molecular replacement)

|  | Ab1303 Fab (PDB 7RYU) | Ab1573 Fab (PDB 7RYV) |
| --- | --- | --- |
| **Data collection** |  |  |
| Space group | P2<sub>1</sub>2<sub>1</sub>2<sub>1</sub> | P2<sub>1</sub> |
| Cell dimensions |  |  |
| *a*, *b*, *c* (Å) | 63.85, 66.97, 136.59 | 81.50 138.40 83.73 |
| $\alpha$, $\beta$, $\gamma$ (°) | 90, 90, 90 | 90 95.71 90 |
| Resolution (Å) | 38.27 - 1.51 (1.53 - 1.51) * | 38.91 – 2.50 (2.78-2.50) * |
| *R*<sub>merge</sub> (%) | 8.4 (124) | 17.4 (98.1) |
| *R*<sub>pim</sub> (%) | 2.4 (37.8) | 10.8 (62.2) |
| CC<sub>1/2</sub> (%) | 99.9 (80.2) | 99.4 (85.8) |
| *I* / $\sigma I$ | 16.5 (1.2) | 8.8 (2.8) |
| Completeness (%) | 99.9 (98.2) | 98.3 (95.9) |
| Multiplicity | 13.3 (11.4) | 3.6 (3.6) |
| **Refinement** |  |  |
| Resolution (Å) | 1.51 | 2.50 |
| No. reflections | 92,598 (9,145) | 63,132 (6,140) |
| *R*<sub>free</sub> / *R*<sub>work</sub> (%) | 20.0 / 18.0 | 25.7 / 22.4 |
| No. atoms |  |  |
| Protein | 3,330 | 12,931 |
| Ligand/ion | 0 | 0 |
| Water | 680 | 880 |
| R.m.s. deviations |  |  |
| Bond lengths (Å) | 0.008 | 0.006 |
| Bond angles (°) | 0.86 | 1.15 |
| Rotamer outliers (%) | 0 | 0.34 |
| Ramachandran plot |  |  |
| Favored (%) | 99.3 | 97.3 |
| Allowed (%) | 0.7 | 2.7 |
| Disallowed | 0 | 0 |
| Average *B*-factor | 27.0 | 30.2 |

\*Values in parentheses are for highest-resolution shell.

**Supplementary table 2.** Cryo-EM data collection, refinement, and validation statistics.

|  | Ab1303-BG505<br>(PDB 7TFN)<br>(EMDB-25877) | Ab1573-BG505<br>(PDB 7TFO)<br>(EMDB-25878) |
| --- | --- | --- |
| **Data collection and processing** |  |  |
| Magnification * | 105,000x | 105,000x |
| Voltage (kV) | 200 | 300 |
| Electron exposure (e-/Å<sup>2</sup>) | 60 | 60 |
| Defocus range (µm) | 1.4-3.0 | 1.4-3.0 |
| Pixel size (Å) | 0.4345 | 0.4275 |
| Recording mode | Super resolution | Super resolution |
| Symmetry imposed | C1 | C1 |
| Initial particle images (no.) | 1,095,744 | 842,259 |
| Final particle images (no.) | 386,825 | 443,817 |
| Overall map resolution (Å) **<br>(masked/unmasked) | 4.0 (4.4) | 4.1 (4.4) |
| **Refinement** |  |  |
| Initial model used (PDB code) | 5CEZ *** | 5CEZ *** |
| Map and model CC | 0.77 | 0.73 |
| Map sharpening *B* factor (Å<sup>2</sup>) | 79.3 | 112 |
| Model composition |  |  |
| Protein residues | 2344 | 2283 |
| Carbohydrate residues | 57 | 21 |
| Validation |  |  |
| MolProbity score | 2.06 | 2.23 |
| Clashscore | 14.1 | 16.4 |
| Poor rotamers (%) | 0 | 0 |
| Ramachandran plot |  |  |
| Favored (%) | 93.9 | 91.1 |
| Allowed (%) | 6.1 | 8.9 |
| Disallowed (%) | 0 | 0 |
| RMS deviations |  |  |
| Length (Å) | 0.003 | 0.005 |
| Angles (°) | 0.63 | 0.70 |

\* Nominal magnification; \*\* FSC threshold 0.143; \*\*\* Partial structure composed of trimeric gp120/gp41

## References

**Uncategorized References**

## References

1. Robertson, D.L., Hahn, B.H. & Sharp, P.M. Recombination in AIDS viruses. Journal of Molecular Evolution 40, 249–259 (1995).

2. Escolano, A., Dosenovic, P. & Nussenzweig, M.C. Progress toward active or passive HIV-1 vaccination. J Exp Med 214, 3–16 (2017).

3. Andrabi, R., Bhiman, J.N. & Burton, D.R. Strategies for a multi-stage neutralizing antibody-based HIV vaccine. Current Opinion in Immunology 53, 143–151 (2018).

4. West, A.P. Jr., et al. Structural Insights on the Role of Antibodies in HIV-1 Vaccine and Therapy. Cell 156, 633–648 (2014).

5. McCoy, L.E. & Burton, D.R. Identification and specificity of broadly neutralizing antibodies against HIV. Immunol Rev 275, 11–20 (2017).

6. Harrison, S.C. Viral membrane fusion. Virology 479-480, 498–507 (2015).

7. Choe, H. et al. The beta-chemokine receptors CCR3 and CCR5 facilitate infection by primary HIV-1 isolates. Cell 85, 1135–48 (1996).

8. Feng, Y., Broder, C.C., Kennedy, P.E. & Berger, E.A. HIV-1 entry cofactor: functional cDNA cloning of a seven-transmembrane, G protein-coupled receptor. Science 272, 872–7 (1996).

9. Sanders, R.W. et al. A next-generation cleaved, soluble HIV-1 Env Trimer, BG505 SOSIP.664 gp140, expresses multiple epitopes for broadly neutralizing but not non-neutralizing antibodies. PLoS Pathog 9, e1003618 (2013).

10. Ward, A.B. & Wilson, I.A. The HIV-1 envelope glycoprotein structure: nailing down a moving target. Immunol Rev 275, 21–32 (2017).

11. Ozorowski, G. et al. Open and closed structures reveal allostery and pliability in the HIV-1 envelope spike. Nature 547, 360–363 (2017).

12. Wang, H., Barnes, C.O., Yang, Z., Nussenzweig, M.C. & Bjorkman, P.J. Partially Open HIV-1 Envelope Structures Exhibit Conformational Changes Relevant for Coreceptor Binding and Fusion. Cell Host Microbe 24, 579–592 e4 (2018).

13. Wang, H. et al. Cryo-EM structure of a CD4-bound open HIV-1 envelope trimer reveals structural rearrangements of the gp120 V1V2 loop. Proc Natl Acad Sci U S A 113, E7151–E7158 (2016).

14. Yang, Z., Wang, H., Liu, A.Z., Gristick, H.B. & Bjorkman, P.J. Asymmetric opening of HIV-1 Env bound to CD4 and a coreceptor-mimicking antibody. Nat Struct Mol Biol 26, 1167–1175 (2019).

15. Burton, D.R. et al. A large array of human monoclonal antibodies to type 1 human immunodeficiency virus from combinatorial libraries of asymptomatic seropositive individuals. Proceedings of the National Academy of Sciences 88, 10134–10137 (1991).

16. Scheid, J.F. et al. Sequence and Structural Convergence of Broad and Potent HIV Antibodies That Mimic CD4 Binding. Science 333, 1633–1637 (2011).

17. Wu, X. et al. Rational design of envelope identifies broadly neutralizing human monoclonal antibodies to HIV-1. Science 329, 856–61 (2010).

18. Zhou, T. et al. Structural definition of a conserved neutralization epitope on HIV-1 gp120. Nature 445, 732–7 (2007).

19. Escolano, A. et al. Sequential immunization elicits broad, but weakly neutralizing, anti-HIV-1 antibodies to the viral envelope V3-glycan patch and CD4bs in rhesus macaques. Submitted (2021).

20. Steichen, J.M. et al. HIV Vaccine Design to Target Germline Precursors of Glycan-Dependent Broadly Neutralizing Antibodies. Immunity 45, 483–96 (2016).

21. Escolano, A. et al. Immunization expands B cells specific to HIV-1 V3 glycan in mice and macaques. Nature 570, 468–473 (2019).

22. McCoy, L.E. et al. Holes in the Glycan Shield of the Native HIV Envelope Are a Target of Trimer-Elicited Neutralizing Antibodies. Cell Rep 16, 2327–38 (2016).

23. Duan, H. et al. Glycan Masking Focuses Immune Responses to the HIV-1 CD4-Binding Site and Enhances Elicitation of VRC01-Class Precursor Antibodies. Immunity 49, 301–311 e5 (2018).

24. Klasse, P.J. et al. Epitopes for neutralizing antibodies induced by HIV-1 envelope glycoprotein BG505 SOSIP trimers in rabbits and macaques. PLoS Pathog 14, e1006913 (2018).

25. Brune, K.D. et al. Plug-and-Display: decoration of Virus-Like Particles via isopeptide bonds for modular immunization. Sci Rep 6, 19234 (2016).

26. Zakeri, B. et al. Peptide tag forming a rapid covalent bond to a protein, through engineering a bacterial adhesin. Proc Natl Acad Sci U S A 109, E690–7 (2012).

27. Wang, Z. et al. Isolation of single HIV-1 Envelope specific B cells and antibody cloning from immunized rhesus macaques. J Immunol Methods 478, 112734 (2020).

28. deCamp, A. et al. Global panel of HIV-1 Env reference strains for standardized assessments of vaccine-elicited neutralizing antibodies. J Virol 88, 2489–507 (2014).

29. Mouquet, H. et al. Complex-type N-glycan recognition by potent broadly neutralizing HIV antibodies. Proc Natl Acad Sci U S A 109, E3268–77 (2012).

30. Walker, L.M. et al. Broad and Potent Neutralizing Antibodies from an African Donor Reveal a New HIV-1 Vaccine Target. Science 326, 285–289 (2009).

31. Kong, R. et al. Fusion peptide of HIV-1 as a site of vulnerability to neutralizing antibody. Science 352, 828–33 (2016).

32. Gristick, H.B. et al. Natively glycosylated HIV-1 Env structure reveals new mode for antibody recognition of the CD4-binding site. Nat Struct Mol Biol 23, 906–915 (2016).

33. Lee, J.H. et al. A Broadly Neutralizing Antibody Targets the Dynamic HIV Envelope Trimer Apex via a Long, Rigidified, and Anionic beta-Hairpin Structure. Immunity 46, 690–702 (2017).

34. Henderson, R. et al. Disruption of the HIV-1 Envelope allosteric network blocks CD4-induced rearrangements. Nature Communications 11(2020).

35. Jette, C.A. et al. Cryo-EM structures of HIV-1 trimer bound to CD4-mimetics BNM-III-170 and M48U1 adopt a CD4-bound open conformation. Nature Communications 12, 1950 (2021).

36. Jeschke, G. DEER distance measurements on proteins. Annu Rev Phys Chem 63, 419–46 (2012).

37. Stadtmueller, B.M. et al. DEER spectroscopy measurements reveal multiple conformations of HIV-1 SOSIP Envelopes that show similarities with Envelopes on native virions. Immunity (2018).

38. Hubbell, W.L., Lopez, C.J., Altenbach, C. & Yang, Z. Technological advances in site-directed spin labeling of proteins. Curr Opin Struct Biol 23, 725–33 (2013).

39. Khramtsov, V.V. et al. Quantitative determination of SH groups in low- and high-molecular-weight compounds by an electron spin resonance method. Anal Biochem 182, 58–63 (1989).

40. Hubbell, W.L., Cafiso, D.S. & Altenbach, C. Identifying conformational changes with site-directed spin labeling. Nature Struct. Biol. 7, 735–739 (2000).

41. Toledo Warshaviak, D., Khramtsov, V.V., Cascio, D., Altenbach, C. & Hubbell, W.L. Structure and dynamics of an imidazoline nitroxide side chain with strongly hindered internal motion in proteins. J Magn Reson 232, 53–61 (2013).

42. Haynes, B.F. & Mascola, J.R. The quest for an antibody-based HIV vaccine. Immunological Reviews 275, 5–10 (2017).

43. Kwong, P.D. et al. Structure of an HIV gp120 envelope glycoprotein in complex with the CD4 receptor and a neutralizing human antibody. Nature 393, 648–59 (1998).

44. Pancera, M. et al. Crystal structures of trimeric HIV envelope with entry inhibitors BMS-378806 and BMS-626529. Nat Chem Biol 13, 1115–1122 (2017).

45. Huang, J. et al. Identification of a CD4-Binding-Site Antibody to HIV that Evolved Near-Pan Neutralization Breadth. Immunity 45, 1108–1121 (2016).

46. Liu, Q. et al. Improvement of antibody functionality by structure-guided paratope engraftment. Nat Commun 10, 721 (2019).

47. Scharf, L. et al. Broadly Neutralizing Antibody 8ANC195 Recognizes Closed and Open States of HIV-1 Env. Cell 162, 1379–90 (2015).

48. Kabsch, W. Integration, scaling, space-group assignment and post-refinement. Acta Crystallogr D Biol Crystallogr 66, 133–44 (2010).

49. Kabsch, W. XDS. Acta Crystallogr D Biol Crystallogr 66, 125–32 (2010).

50. Winn, M.D. et al. Overview of the CCP4 suite and current developments. Acta Crystallogr D Biol Crystallogr 67, 235–42 (2011).

51. McCoy, A.J. et al. Phaser crystallographic software. J Appl Crystallogr 40, 658–674 (2007).

52. Adams, P.D. et al. PHENIX: a comprehensive Python-based system for macromolecular structure solution. Acta Crystallogr D Biol Crystallogr 66, 213–21 (2010).

53. Afonine, P.V. et al. Real-space refinement in PHENIX for cryo-EM and crystallography. Acta Crystallogr D Struct Biol 74, 531–544 (2018).

54. Emsley, P., Lohkamp, B., Scott, W.G. & Cowtan, K. Features and development of Coot. Acta Crystallogr D Biol Crystallogr 66, 486–501 (2010).

55. Zheng, S.Q. et al. MotionCor2: anisotropic correction of beam-induced motion for improved cryo-electron microscopy. Nat Methods 14, 331–332 (2017).

56. Rohou, A. & Grigorieff, N. CTFFIND4: Fast and accurate defocus estimation from electron micrographs. J Struct Biol 192, 216–21 (2015).

57. Zivanov, J. et al. New tools for automated high-resolution cryo-EM structure determination in RELION-3. Elife 7(2018).

58. Scheres, S.H. RELION: implementation of a Bayesian approach to cryo-EM structure determination. J Struct Biol 180, 519–30 (2012).

59. Tan, Y.Z. et al. Addressing preferred specimen orientation in single-particle cryo-EM through tilting. Nature Methods 14, 793–796 (2017).

60. Lefranc, M.P. et al. IMGT(R), the international ImMunoGeneTics information system(R) 25 years on. Nucleic Acids Res 43, D413–22 (2015).

61. Goddard, T.D. et al. UCSF ChimeraX: Meeting modern challenges in visualization and analysis. Protein Sci 27, 14–25 (2018).

62. Krissinel, E. & Henrick, K. Inference of macromolecular assemblies from crystalline state. J Mol Biol 372, 774–97 (2007).

63. Dolinsky, T.J. et al. PDB2PQR: expanding and upgrading automated preparation of biomolecular structures for molecular simulations. Nucleic Acids Research 35, W522–W525 (2007).

64. Baker, H.M., Mason, A.B., He, Q.Y., MacGillivray, R.T. & Baker, E.N. Ligand variation in the transferrin family: the crystal structure of the H249Q mutant of the human transferrin N-lobe as a model for iron binding in insect transferrins. Biochemistry 40, 11670–5 (2001).

65. Fleissner, M.R. et al. Site-directed spin labeling of a genetically encoded unnatural amino acid. Proc Natl Acad Sci U S A 106, 21637–42 (2009).

